# Socially regulated genes are spatially hyperconnected to enhancers in the ant brain

**DOI:** 10.64898/2026.04.08.717219

**Authors:** Mohuizi Kuang, Aneliya Antonova, Sandra Moreno-Medina, John F. Doherty, Kavitha Sarma, Marco Prinz, H.T. Marc Timmers, Emily J. Shields, Roberto Bonasio

## Abstract

Caste identity in *Harpegnathos saltator* ants remains plastic beyond development and throughout adulthood. Adult *Harpegnathos* workers can become dominant reproductives, known as “gamergates,” through a social caste transition that involves extensive transcriptional and cellular remodeling of the brain. To gain insight into the epigenetic regulation of this process, we generated comprehensive, caste-specific epigenomic atlases of the *Harpegnathos* brain, including chromatin accessibility, histone modifications, and 3D genome architecture. Using these data we refined the genome assembly, annotated enhancers, and linked them to target genes. We then identified candidate 3D-architectural factors, many of which were specifically upregulated in gamergate brains. Promoters of genes upregulated during the worker–gamergate transition formed an unusually high number of 3D chromatin contacts with their regulatory regions, and most of these contacts were already present in workers. We propose that the pre-existing hyper-connectivity of socially regulated genes is essential to adult brain plasticity and behavioral reprogramming.

## INTRODUCTION

Phenotypic plasticity manifests itself in cells, tissues, and even entire organisms as they respond to developmental cues and environmental changes (Borrelli et al., 2008; Merrell and Stanger, 2016; Sultan, 2021). Plasticity is central to the success of eusocial insects, allowing most species to define reproductive queen or non-reproductive worker castes during the larval stage, thereby establishing division of labor and social organization within colonies (Corona et al., 2016). In some more evolutionarily ancestral ant species, including *Harpegnathos saltator* (henceforth *Harpegnathos*), individuals retain remarkable behavioral and brain plasticity even in adulthood (Peeters, 1991; Peeters et al., 1997). In the absence of a dominant reproductive, adult *Harpegnathos* workers duel among themselves and some transition into long-lived, reproductive pseudo-queens, termed “gamergates” (Peeters and Holldobler, 1995). Established gamergates can also revert back to worker status following change in the social context (Penick et al., 2021; Yan et al., 2022). Through the worker–gamergate transition, individuals gain reproductive ability, social dominance, ∼5-fold longer lifespan, and a new behavioral repertoire (Ghaninia et al., 2017; Peeters et al., 2000). These phenotypic and behavioral changes are accompanied by dramatic alterations in gene expression, hormone levels, and cellular composition in the brain (Gospocic et al., 2021; Gospocic et al., 2017; Opachaloemphan et al., 2021; Sheng et al., 2020; Yan et al., 2022).

Several lines of evidence demonstrate that epigenetic pathways underpin healthy brain function, plasticity, and disease in humans and model organisms (Bonasio, 2012; Borrelli et al., 2008; Campbell and Wood, 2019; Frost et al., 2014; Gallegos et al., 2018; Griffith et al., 2024; Hwang et al., 2017; Nativio et al., 2018; Robison and Nestler, 2011). The profound reprogramming of the adult brain in *Harpegnathos* ants offers a unique model to study chromatin and gene regulation in a relatively simple, yet eminently plastic brain. Studies in other eusocial insects have already demonstrated that histone marks, such as H3 lysine 27 acetylation (H3K27ac), are linked to caste-specific gene expression and behavior (Simola et al., 2013). Transcription factors (TFs) are also recognized as drivers of brain plasticity, not only in *Harpegnathos* (Gospocic et al., 2021), but also in more complex mammalian brains (Alberini, 2009; McClard et al., 2018).

*Cis*-regulatory elements (CREs) including promoters, enhancers, and insulators are bound by TFs, which regulate gene expression by recruiting chromatin-modifying enzymes, remodelers and the transcriptional machinery. This mechanism is in part enabled by three-dimensional (3D) chromatin folding, which allows CREs to come in spatial proximity to their target genes to coordinate gene expression (Yang and Hansen, 2024). In the brain, genome folding orchestrates neuronal function by establishing cell-type-specific transcriptional programs essential for neuronal identity and circuit formation (Bonev et al., 2017; Fujita et al., 2022; Marco et al., 2020; Rahman et al., 2023; Wahl et al., 2024). 3D chromatin contact maps in the human brain have identified promoters that display hyperconnectivity, termed “super interactive promoters” (SIPs), which drive expression of lineage-specific genes and act as critical nodes to maintain cell-type identity (Heffel et al., 2024; Song et al., 2020). Despite these insights, whether and how 3D genome organization contributes to brain plasticity remains largely unknown.

To address this gap, we constructed a comprehensive regulatory atlas of *Harpegnathos* brains by generating a multi-layered dataset profiling the transcriptome, linear chromatin landscape, and 3D genome organization. We utilized the dataset to first refine genome assembly and annotate putative enhancers. Next, we characterized patterns of regulatory architecture based on chromatin features and chromatin looping. Finally, we inspected chromatin folding at caste-specific genes and found an unexpectedly high level of connectivity at promoters of genes upregulated during the social transition.

## RESULTS

### Epigenomic profiling and chromatin states of the *Harpegnathos* brain

To characterize the chromatin structure of the *Harpegnathos* brain, we constructed a comprehensive atlas of the regulatory genome covering the linear chromatin landscape (accessibility and histone marks), transcriptome, and 3D chromatin architecture. We optimized ATAC-seq and CUT&Tag assays for ant brains to profile genome-wide chromatin accessibility, as well as a panel of well-studied histone modifications to define epigenetic states and classes of regulatory elements. Specifically, we chose histone marks typically associated with active promoters (H3K4me3), active enhancers (H3K4me1, H3K27ac), transcribed gene bodies (H3K36me3), *Polycomb*-repressed chromatin (H2AK119ub, H3K27me3), and constitutive heterochromatin (H3K9me3) (Roadmap Epigenomics et al., 2015). For each mark, we generated multiple biological replicates from young workers (34–43 days post-eclosion) by harvesting their brain and removing the optic lobes (henceforth “non-visual brain”; **Fig. 1A**).

**Figure 1.**
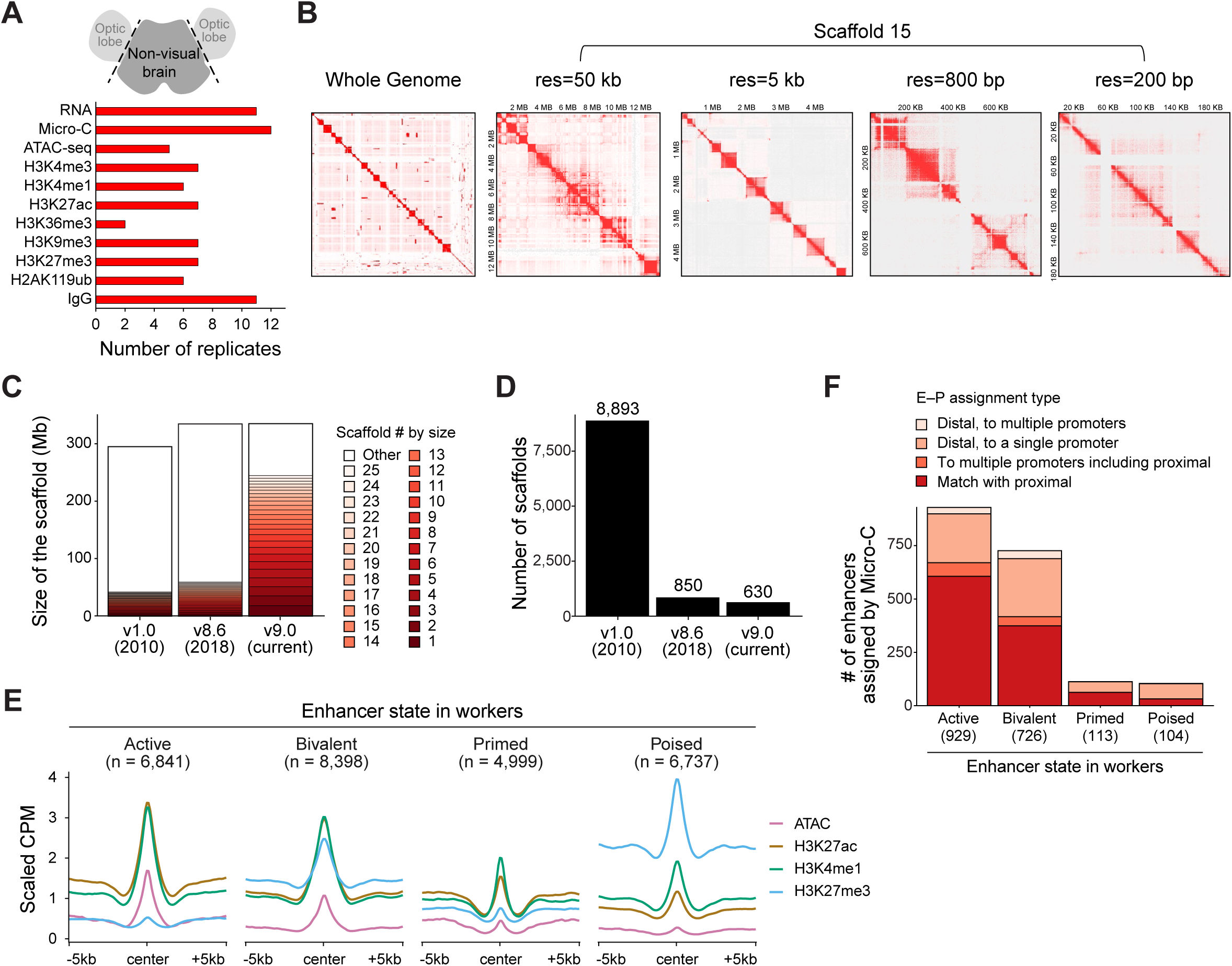
3D chromatin maps improve genome assembly and enhancer annotation. (A) Top, illustration of the part of the brain used for all experiments, labeled “non-visual brain”. Bottom, number of biological replicates of brains profiled using RNA-seq, Micro-C, ATAC-seq, or CUT&Tag for H3K4me3, H3K4me1, H3K27ac, H3K36me3, H3K9me3, H3K27me3, H2AK119ub, or for IgG control. (B) Micro-C contact maps for the whole-genome scale (left) and for portions of scaffold 15 (*Hsal* 9.0) at increasing resolutions (res). The matrices are shown as KR-normalized, balanced contact frequencies. (C) Size in millions of base pairs for the longest 25 scaffolds (red colors) in comparison to all other scaffolds (white) in assembly version v1.0 (Bonasio et al., 2010), v8.6 (Shields et al., 2018), and v9.0 (this study). (D) Number of scaffolds in assembly version v1.0 (Bonasio et al., 2010), v8.6 (Shields et al., 2018), and v9.0 (current version). (E) ATAC, H3K27ac, H3K4me1, and H3K27me3 signal at enhancers (center ± 5 kb). Enhancers were classified into active, bivalent, primed, and poised based on their accessibility and histone marks in worker brains. The *y* axis represents raw counts per million (CPM) for ATAC, or scaled, IgG-normalized CPM (for CUT&Tag, see methods). (F) Comparison of assignments of enhancers to target genes using Micro-C and with proximity assignment method. Numbers of active, bivalent, primed, and poised enhancers (as defined by their chromatin state in worker brains) for which gene target assignment using Micro-C and proximity coincided (“match with proximal”), matched to multiple promoters including the proximal promoter, or matched to distal promoters. The number of biological replicates for all analyses is shown in panel (A).

We confirmed the validity of our chromatin profiles by examining the genomic localization of each mark. Compared to the background genome, H3K4me3 and ATAC-seq peaks were enriched at promoters; H3K36me3 was preferentially located at exons; H3K27ac, H3K4me1, H3K27me3, and H2AK119ub were distributed over all features, with a slight enrichment in introns; and H3K9me3 was primarily found in intergenic regions (**Fig. S1A**). To examine the relationship between the chromatin landscape and gene expression, we also acquired 11 individual RNA-seq profiles from worker brains (**Fig. 1A**) and observed the expected correlations between transcriptional activity and chromatin features (**Fig. S1B**). Specifically, the promoters of highly expressed genes were accessible, as evidenced by high ATAC-seq signal, and were enriched for H3K4me3 and H3K27ac, while their gene bodies displayed robust H3K36me3 deposition. Conversely, lowly expressed genes were enriched in histone marks typically associated with repressed heterochromatin, namely H2AK119ub, H3K27me3, and H3K9me3.

We utilized ChromHMM analysis (Ernst and Kellis, 2017) to partition the genome into 15 chromatin states defined by distinctive combinatorial patterns of chromatin features and their known functional associations (**Fig. S1C**). Active/flanking transcription start sites (a/flTSS) and active gene bodies (aGB) were defined by marks associated with active transcription, such as H3K4me3 and H3K36me3, while Polycomb-repressed (RepP) and heterochromatic (Het) states were characterized by H3K27me3 (with or without H2AK119ub) and H3K9me3, respectively. These ChromHMM annotations were validated by correlation with transcriptional activity: genes with promoters in active states were highly expressed, whereas genes with promoters in repressed states exhibited low expression (**Fig. S1D**).

To our knowledge, this dataset represents the most comprehensive panel of chromatin accessibility and histone modification profiles in a social insect brain. It displays the expected correlations between gene expression and chromatin features, and enabled the identification of functional and regulatory elements in the *Harpegnathos* genome, as described below.

### Refined genome assembly and enhancer annotation through 3D chromatin contact maps

To determine the 3D chromatin organization of the 334 Mb *Harpegnathos* genome, we performed Micro-C (Hsieh et al., 2015) to generate single-nucleosome resolution 3D contact maps. We adapted the Micro-C protocol (Slobodyanyuk et al., 2022) for low-input samples and collected 12 biological replicates across two batches, each biological replicate consisting of an individual worker’s non-visual brain (**Fig. 1A**). Deep sequencing of these libraries produced approximately 1.04 billion total reads (**Fig. S2A–B**, **Table S1**). The resulting high-resolution contact maps enabled us to visualize canonical features of 3D chromatin architecture, including chromosome territories, A/B compartments, and topologically associated domains (TADs) across different scales (**Fig. 1B**).

Previously, we used short-read (*Hsal* v1.0) (Bonasio et al., 2010) and PacBio long-read (*Hsal* v8.6) (Shields et al., 2018) to assemble the *Harpegnathos* genome, but even in our latest, higher-quality assembly (*Hsal* v8.6), the genome remained fragmented into 850 scaffolds. Given that intra-chromosomal contacts are much more frequent than inter-chromosomal interactions, genome-wide 3D contact maps (e.g., derived from Hi-C or Micro-C) can be used to connect physically adjacent chromosome regions, therefore improving the continuity of genome assemblies (Burton et al., 2013; Dudchenko et al., 2017; Kim et al., 2025). Using a set of 164 million Micro-C reads aimed at improving the assembly, separate from the larger set described above (see methods), we connected disjointed scaffolds from the 2018 *Hsal* v8.6 assembly, quadrupling the genome size covered by the 25 largest scaffolds (**Fig. 1C**) and reducing the total scaffold number from 850 (v8.6) to 630 (*Hsal* v9.0, current assembly) (**Fig. 1D**). The N50 size, a conventional metric for genome assembly quality, increased by ∼8-fold from 1.08 M to 8.4 M. As an example, the *msi* locus, which was split into two scaffolds in the *Hsal* v8.6 assembly, was connected in the *Hsal* v9.0 assembly (**Fig. S2C**). This new and refined assembly was used for all subsequent analyses.

As distal CREs, enhancers serve as docking sites for TFs, which can bring them in the physical vicinity of their target genes via chromatin looping (Kim and Wysocka, 2023). We annotated 26,973 putative enhancers in the *Harpegnathos* brain based on characteristic chromatin signatures (Heintzman et al., 2007) from the worker profiles and adding 6 replicates of H3K4me1 CUT&Tag from age-matched gamergate brains, to capture potential caste-specific enhancers. We next classified enhancers into active, bivalent, primed, or poised, based on the overlap with specific chromatin signatures in worker brains, including accessibility (ATAC-seq), H3K27ac, and/or H3K27me3 (Heinz et al., 2015). As a result, we defined 6,841 active enhancers (accessible or marked by H3K27ac and devoid of H3K27me3), 8,398 bivalent enhancers (accessible or marked by H3K27ac and also carrying the H3K27me3 modification), 4,999 primed enhancers (–ATAC, –H3K27ac, –H3K27me3), and 6,737 poised enhancers (–ATAC, –H3K27ac, +H3K27me3) (**Fig. 1E**).

In the absence of 3D conformation information, standard bioinformatic approaches typically rely on “the nearest gene” rule for linking putative enhancers to their linearly immediate proximal genes; however, the method overlooks instances where an enhancer “skips” intervening genes to target a distal promoter, and can result in misannotations (Kim and Wysocka, 2023; Shen et al., 2012). We therefore utilized chromatin loops derived from Micro-C data to assign enhancers to genes through identified enhancer–promoter (E–P) contacts. To maximize chances to detect caste-specific contacts, we obtained a comparable dataset of Micro-C from gamergate brains (**Table S2**).

Among all annotated enhancers (26,973), 1,872 (∼7%) formed detectable chromatin loops with one or more promoters (**Fig. 1F**). As expected, the majority of the promoter-contacting enhancers were active (929, 12.93% of active enhancers), or bivalent (726, 8.4% of bivalent enhancers) in worker brains; whereas primed enhancers (113, 2.4%) and poised enhancers (104, 1.5%) were less likely to form a detectable contact with promoters (**Fig. 1F**). While in many cases, loops connected enhancers to the most proximal promoter, a substantial number of enhancers (690, ∼37% of enhancers with detectable E–P loops) skipped the adjacent gene and contacted single or multiple distal promoters (**Fig. 1F**, and see example in **Fig. S2D**). Genes assigned to active enhancers by Micro-C were enriched for gene ontology (GO) terms related to neuronal and brain function including “*signal transduction*,” “*calcium ion transmembrane transport*,” and “*chemical synaptic transmission*” (**Fig. S2E**).

Collectively, these 3D chromatin contact maps allowed us to improve the continuity of the *Harpegnathos* genome assembly. By combining Micro-C data with histone modification profiles, we annotated putative enhancers, predicted their functional states, and linked them to target genes. These resources laid the foundations for our subsequent analyses of genome organization and chromatin regulation in the *Harpegnathos* brain.

### Active promoters form TAD boundaries

Topologically associating domains (TADs) are structural and functional units of 3D genome organization, comprised of stretches of chromatin that preferentially self-interact within their boundaries and are largely insulated from interactions with neighboring domains (Dixon et al., 2012; Hou et al., 2012; Nora et al., 2012; Sexton et al., 2012). TAD boundaries are therefore considered to be architectural insulators, essential for guarding proper gene regulation within their respective neighborhoods. We identified 4,608 TADs in the *Harpegnathos* brain and marked TAD boundaries by calculating genome-wide insulation scores on our Micro-C data at different resolution sizes (**Fig. 2A**, “insulation”). For subsequent TAD analyses, we chose 400 bp resolution with 10 kb windows, which balanced detection sensitivity and smoothness of the resulting insulation profiles. TAD boundaries were highly reproducible across individuals, as demonstrated by the consistency among the insulation profiles of the 12 biological replicates across two experimental batches (**Fig. S3A–B**).

**Figure 2.**
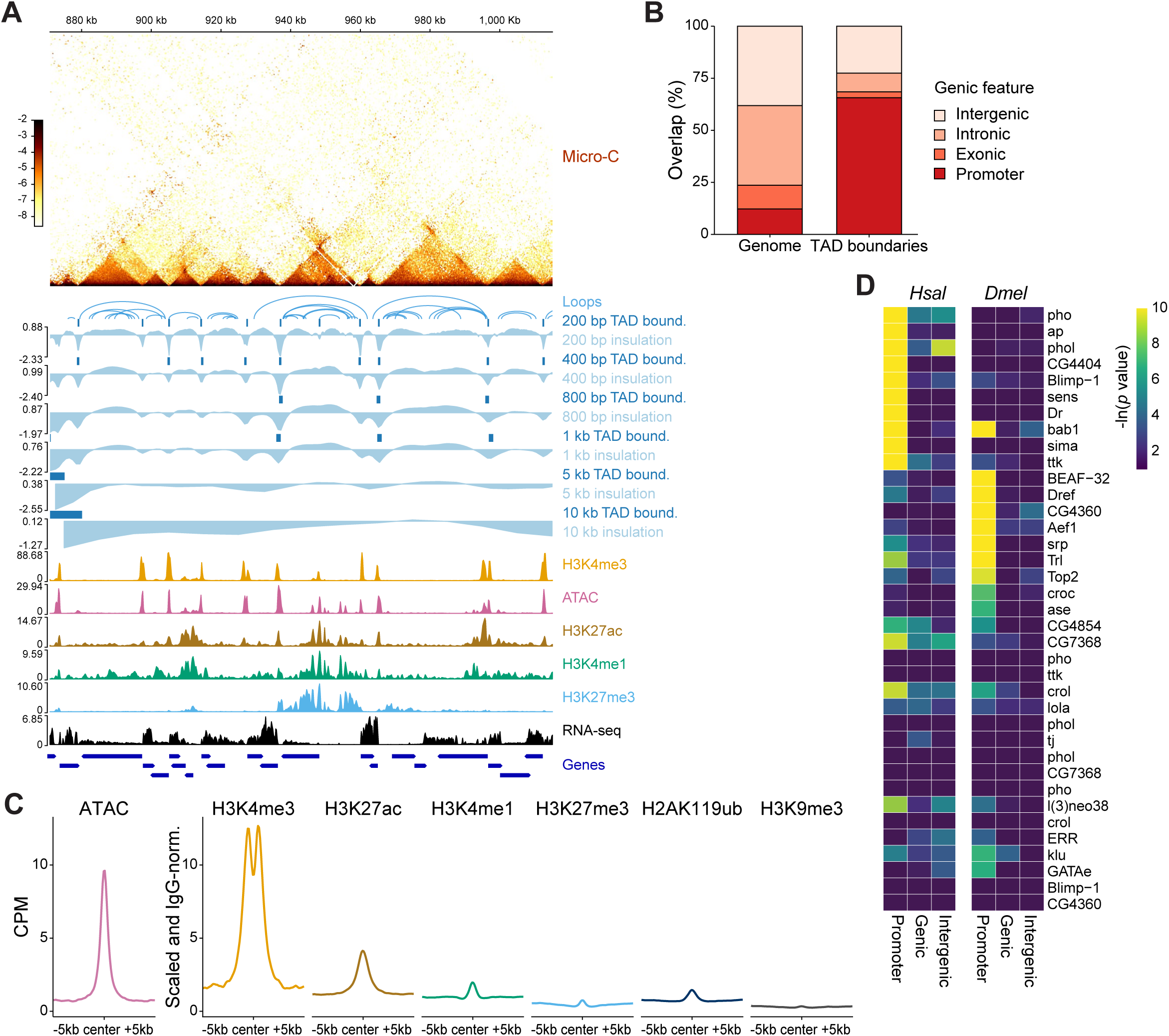
TAD boundaries are enriched at active promoters in *Harpegnathos*. (A) Micro-C and chromatin enrichment across a 1 Mb region on scaffold 49. Chromatin interactions are visualized as balanced, Knight-Ruiz (KR)-normalized contact frequencies. TAD boundaries at different resolutions were identified by insulation score. RNA-seq, ATAC-seq and CUT&Tag signal for H3K4me3, H3K27ac, H3K4me1, H3K27me3 are shown in CPM along with annotated genes. (B) Percentage of TAD boundaries overlapping with promoter (± 1 kb from the TSS), exonic, intronic, and intergenic regions. (C) ATAC-seq (CPM) and CUT&Tag signal (H3K4me3, H3K27ac, H3K4me1, H3K27me3, H2AK119ub, and H3K9me3, normalized with scaled IgG) enrichment at all TAD boundaries. (D) Motif enrichment at TAD boundaries overlapping with promoter, genic, or intergenic regions in *Harpegnathos* or *Drosophila*, identified by SEA-based analysis using shuffled sequences preserving dinucleotide frequencies as the background.

In mammals and *Drosophila*, TAD boundaries are enriched for the active promoter mark H3K4me3, housekeeping genes, and transfer RNAs (Dixon et al., 2012; Sexton et al., 2012). Consistent with these findings, in *Harpegnathos*, TAD boundaries were predominantly localized to promoter regions rather than gene bodies or intergenic regions (**Fig. 2A–B**). Genic TAD boundaries were enriched for 5’ UTR and tRNA genes, while being depleted of transposable elements such as LINEs (**Fig. S3C**). Promoter enrichment at TAD boundaries was further highlighted by metaplots revealing strong signals for chromatin accessibility, active promoter mark H3K4me3, and a moderate enrichment of H3K27ac, which were all localized at the boundary center (**Fig. 2C**). To quantify the association between boundary strength and chromatin states, we divided TAD boundaries according to their ChromHMM states and calculated the average boundary strength linked to each state. TAD boundaries at active TSS acted as the strongest insulators (**Fig. S3D**), indicating that promoters, especially active promoters, formed the most frequent and strongest obstacle to chromatin interactions across TADs.

In vertebrates, TAD boundaries are characterized by the co-localization of the CCCTC-binding factor (CTCF) and the structural maintenance of chromosomes (SMC) cohesin complex, whereas in *Drosophila*, they are bound by a more diverse repertoire of TFs with architectural functions, such as BEAF-32, Su(Hw), CP190, and Dref (Acemel and Lupianez, 2023; Dixon et al., 2012; Gurudatta et al., 2013; Ramirez et al., 2018). We searched for DNA sequences enriched at TAD boundaries in *Harpegnathos* using a database of motifs bound by *Drosophila* TFs and, as a comparison, we performed the same analysis on Micro-C from adult *Drosophila* brains (Mohana et al., 2023). As in *Harpegnathos* (**Fig. 2B**), *Drosophila* TAD boundaries were predominantly located at promoter regions (**Fig. S3E**) and were enriched for motifs recognized by known architectural proteins, including BEAF-32 and Dref (**Fig. 2D**). Some of the motifs found at *Harpegnathos* boundaries were shared with *Drosophila* (e.g. those recognized by bab1 and crol; **Fig. 2D**), suggesting an evolutionarily conserved role for these factors in orchestrating 3D genome architecture in insect brains. In contrast, motifs for other TFs (e.g. sima and Dr), were only enriched in *Harpegnathos* TAD boundaries, suggesting evolutionary divergence in genome architecture principles.

Together, our results highlight both conserved and distinct characteristics of TAD insulation in the *Harpegnathos* brain: TAD boundaries frequently coincided with active promoters, which is a feature conserved across diverse species. Some of the TFs delineating TAD boundaries have a conserved architectural role in *Drosophila*, whereas others are unique to *Harpegnathos*, and might play similar roles in other social insects.

### Genetic and epigenetic features of TAD, loops, and meta-loops

We annotated six classes of TADs in *Harpegnathos* brains, based on their histone modification signatures and inferred epigenetic states (**Fig. 3A, S4A**): two classes of “active” TADs, Polycomb-repressed TADs (“PcG”), H3K9me3-marked heterochromatic TADs (“Het”); a “bivalent” class marked by moderate H3K27ac, H3K4me1, H3K27me3, and H2AK119ub; and a set of “undefined” TADs with low signal for multiple histone marks, likely reflecting cellular heterogeneity or experimental noise. Overall, 20% of the *Harpegnathos* genome was covered by active TADs, which were the smallest in size, while repressive TADs spanned 28% of the genome and were in average larger (**Fig. 3B–C**).

**Figure 3.**
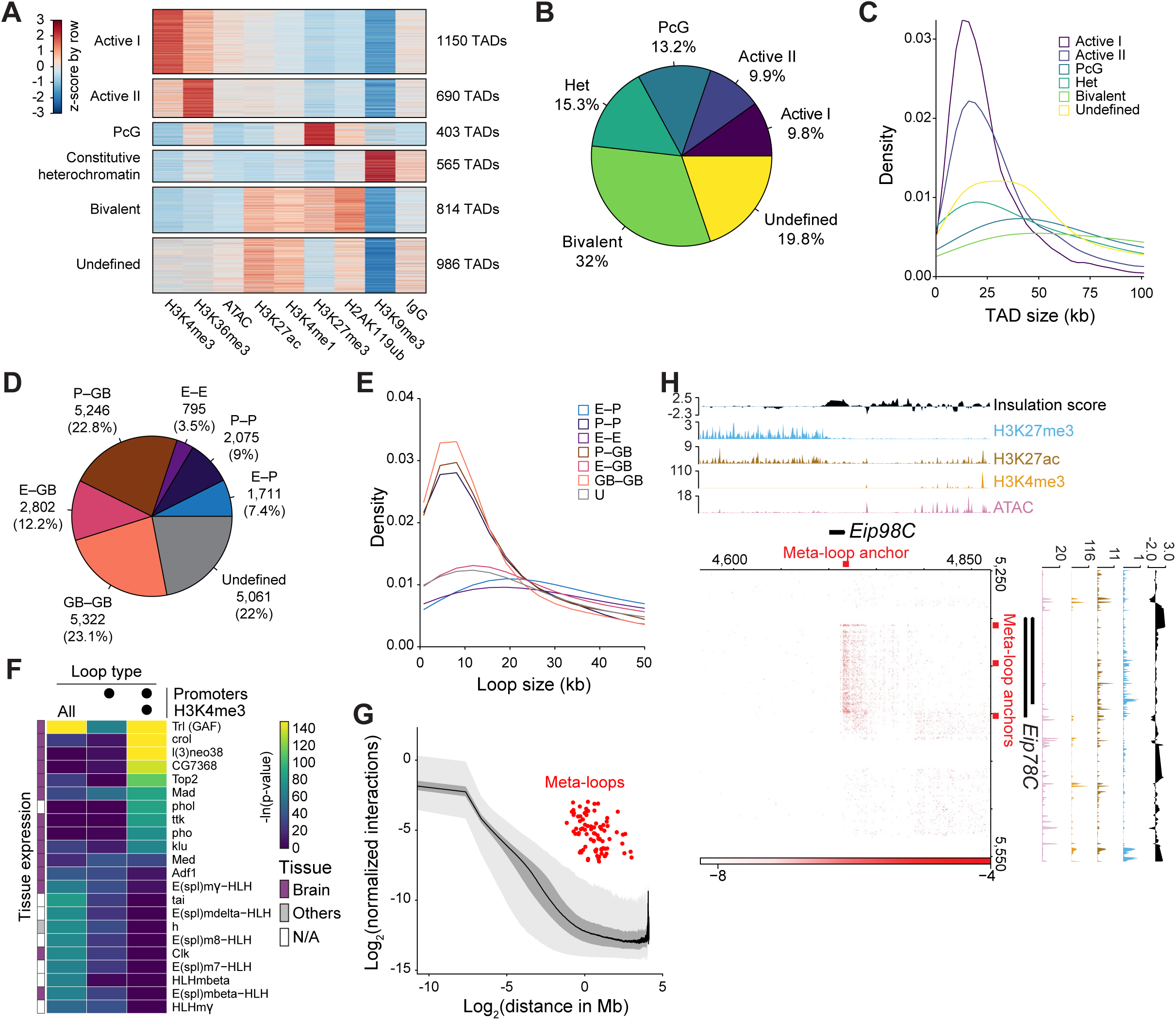
Genetic and epigenetic features of TADs, loops, and meta-loops. (A) K-means clustering of 4,608 TADs into 6 classes, annotated based on their chromatin enrichment. Each row represents a TAD. (B) Percentage of the genome covered by each TAD class. (C) Size distribution of each TAD class. (D) Percentage of chromatin loops classified as promoter–promoter (P–P), enhancer–promoter (E–P), promoter–gene body (P–GB), enhancer–gene body (E–GB), gene body–gene body (GB–GB), or “undefined”, indicating that at least one of the two anchors maps to an intergenic region devoid of enhancers. (E) Distribution of anchor distance in kb for each class of chromatin loop. (F) Motif enrichment at all loop anchors against a shuffled background (left column), at all promoters with loops against promoters without loops (middle column), or promoters with loops and marked by H3K4me3 against promoters with loops but devoid of H3K4me3 (right column). Preferential tissue expression is indicated on the left (see also Fig. S5C). (G) Distribution of genome-wide normalized interaction frequency against distance. Red dots show identified meta-loops; light and dark gray curves show 1st–99th or 25th–75th percentile ranges of all contact pairs, respectively; black line shows the median. (H) An example meta-domain formed by two TADs comprising *Eip98C* (horizontal) and *Eip78C* (vertical) connected by three meta-loops, anchored at sites indicated by red blocks at the margins of the interaction matrix. The Micro-C contact frequencies, KR-normalized and balanced are shown in the center matrix; insulation score, ATAC-seq, and CUT&Tag signal for H3K4me3, H3K27me3, H3K27ac are plotted along the margins for the two loci.

Both repressive TAD types were often adjacent to TADs of the same class (**Fig. S4B**), and boundary strengths were the highest for PcG TADs (**Fig. S4C**). Both active TAD types preferentially comprised expressed genes and active promoters (**Fig. S4D–E**), whereas active II TADs preferentially contained active enhancers (**Fig. S4F**). PcG TADs comprised a high density of poised or bivalent enhancers (**Fig. S4F**), as expected for developmentally controlled regions of heterochromatin, whereas Het TADs were enriched for repetitive elements, consistent with the constitutive heterochromatin state (**Fig. S4G**). PcG TADs contained genes involved in neuronal and cellular differentiation, as well as chemosensory perception, suggesting that the Polycomb pathway silenced these genes in the adult central nervous system (Sieriebriennikov et al., 2025; Yan et al., 2020). On the other hand, genes involved in “synaptic organization” were found within bivalent TADs, likely indicating heterogeneous gene expression patterns in the different cell types in the brain (**Fig. S4H**).

We identified 23,012 chromatin loops in the *Harpegnathos* brain, of which at least 4,581 (∼20%) connected two annotated CREs (promoters or enhancers), including 1,711 E–P loops (**Fig. 3D**). Compared to all other loops, loops anchored at enhancers, including E–P, enhancer–enhancer (E–E), and enhancer–gene body (E–GB) loops, spanned in average greater genomic distances, with median lengths of 42 kb (**Fig. 3E**), highlighting long-distance enhancer-based gene regulation in the *Harpegnathos* brain.

Compared with *Drosophila*, *Harpegnathos* brains displayed a relative expansion of enhancer-anchored loops (23.1% versus 14.9%) (**Fig. S5A**, thick outlines). Promoters contacting active enhancers were linked to genes with the highest RNA levels (**Fig. S5B**), confirming the functional relevance of enhancer-associated loops. The motif recognized by the pioneer TF GAGA-associated factor (GAF), encoded by the *Trl* gene, was greatly enriched at all loop anchors (**Fig. 3F**, left column), which is consistent with its known role in 3D genome organization in *Drosophila* (Li et al., 2023). The motif recognized by GAF/Trl persisted was also strongly enriched in loop anchors found at promoters (**Fig. 3F**, middle column), as well as in loop anchors at active promoters (**Fig. 3F**, right column). In the latter class of loops, we found additional enriched motifs including those for TFs encoded by *ttk*, *Mad,* and *pho*, suggesting a specific role for these factors in 3D genome organization at active genes in the ant brain. In fact, a large majority of these TFs were preferentially expressed in the *Harpegnathos* brain, compared to other tissues (Shields et al., 2018) (**Fig. 3F**, **Fig. S5C**). Some of the same motifs were also enriched at loop anchors in *Drosophila* adult brains (**Fig. S5D**), suggesting a potential conservation of the architectural function of the corresponding TFs across > 300 million years of evolutionary divergence.

A recent study of genome organization in the *Drosophila* brain reported interactions between distant TADs mediated by ultra-long-distance “meta-loops”, which facilitate co-regulation of loci far apart in sequence space (Mohana et al., 2023). We identified 98 inter-TAD meta-loops in the *Harpegnathos* brain, spanning distances from 445 kb to 8 Mb (**Fig. 3G**) and primarily anchored within active TADs (**Fig. S5E**). Similar to a *Drosophila* meta-loop connecting the glutamate receptor IA and IB paralogs (*GluRIA* and *GluRIB*) (Mohana et al., 2023), multiple meta-loops connected two TADs comprising ecdysone-induced genes *Eip78C* and *Eip98C* (**Fig. 3H**), which reside 687 kb apart and are simultaneously upregulated during the worker–gamergate transition (Gospocic et al., 2021). This suggests that 3D-based mechanisms to co-regulate genes acting in the same pathway in the brain might be conserved from *Drosophila* to ants.

In summary, 3D genome organization in the *Harpegnathos* brain presents several conserved features compared with *Drosophila*, including brain-specific TFs that potentially serve as chromatin looping factors, and meta-loops that might be deployed to coordinate gene activation during the social caste transition. We also found that E–P interactions are more prominent in the ant brain compared to the fly brain, motivating further analyses on enhancer and promoter connectivity in *Harpegnathos*.

### Motifs for gamergate TFs are enriched at socially regulated loop anchors

*Harpegnathos* adult ants retain profound brain and behavioral plasticity, allowing workers to transition into reproductives called gamergates (Carmona-Aldana et al., 2024; Libbrecht et al., 2013; Yan et al., 2014). Nothing is known about how epigenetic and 3D remodeling of the genome contribute to this remarkable example of adult brain plasticity. We reprogrammed workers to gamergates by separating 20–25 callows from their colony of origin and allowing them to re-establish a social hierarchy over 30 days (**Fig. 4A**). After dueling, individuals were classified into workers or gamergates based on behavioral observations and ovary scoring, which clearly distinguished the two behavioral castes (**Fig. S6A**). Transcriptome analyses revealed 1,660 differentially expressed genes, including well-known caste-biased genes, such as corazonin (*Crz*), neuroparsin-A (*NpA*), and Krüppel homolog-1 (*Kr-h1*) (Gilbert et al., 2025; Gospocic et al., 2021; Gospocic et al., 2017; Opachaloemphan et al., 2021), which were preferentially expressed in the worker brain, whereas vitellogenin (*Vg*) and juvenile hormone acid O-methyltransferase 3 (*jhamt3*) (Gospocic et al., 2021; Gospocic et al., 2017; Yan et al., 2022) were upregulated in the gamergate brain (**Fig. 4B**, **S6B**). Aligning with our previous observations (Sheng et al., 2020), our dataset confirmed that the transcriptional programs of workers and gamergates after 30 days of transition were largely finalized, as they were highly correlated with those of mature workers and gamergates after 120 days of transition (**Fig. S6C**) (Gospocic et al., 2021).

**Figure 4.**
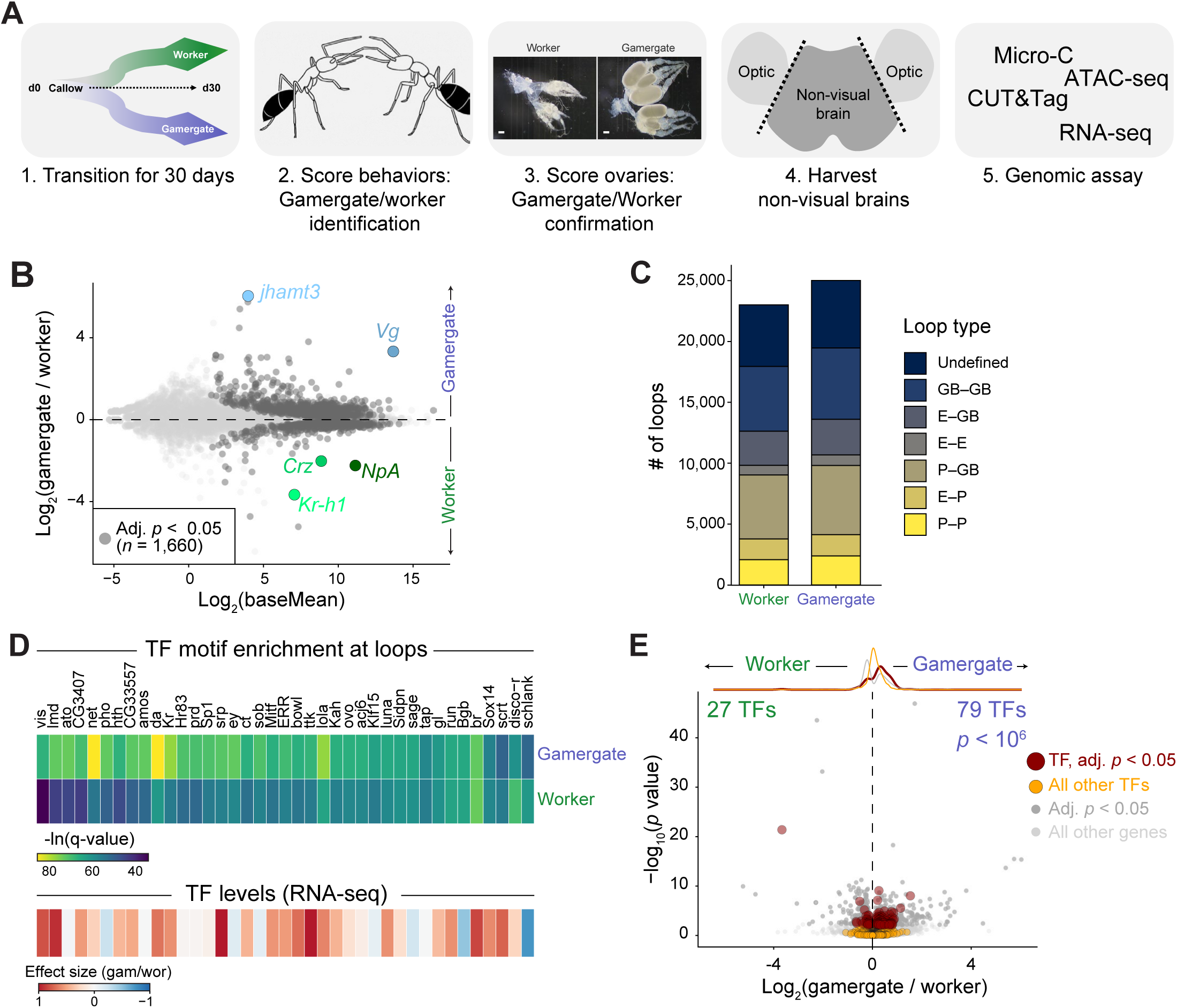
Motifs for gamergate TFs are enriched at gamergate loop anchors. (A) Experimental scheme. Young workers were isolated from stable colonies and placed in a new empty nest, where they engaged in dueling tournaments leading to the transition of some individuals to reproductive gamergates (1). Workers and gamergates were identified through behavioral scoring (2) and their identities were confirmed after harvesting by inspecting their ovaries (3). Non-visual brains were harvested at 30 days after isolation (4) and used for various genomic assays. (B) MA plot of RNA-seq data from day-30 gamergates (*n* = 17) and workers (*n* = 11). Significantly differentially expressed genes are shown in dark gray and known caste-specific genes are highlighted. (C) Number of each chromatin loop type detected in worker and gamergate brains. (D) Motif enrichment at anchors of worker– and gamergate-specific differential loops using shared loop anchors as the background, with scale showing –log(q-value) (top). Also shown is RNA-seq effect size, defined as log_2_(gamergate/worker) / standard error, of the corresponding TFs. (E) RNA-seq data from (B) with non-differentially expressed TFs (orange) and differentially expressed TFs (red) highlighted. The histograms above the volcano plot show the distributions of each set of genes. *P* value is from a Fisher’s exact test for the proportion of TFs upregulated in each caste.

Next, we compared 3D chromatin conformation in worker and gamergate brains. Despite comparable sequencing depths (**Table S1–2**), chromatin in the gamergate brain formed 8.7% more loops (25,004) than in the worker brain (23,012) (**Fig. 4C**). To uncover a potential regulatory basis for looping differences, we searched for TF motifs enriched at anchors of caste-specific loops vs. anchors shared between castes. We found a set of caste-biased TF motifs, the majority of which were preferentially localized at gamergate-specific loop anchors (**Fig. 4D**, top). Most of the corresponding TFs (29/37, 78%) were also expressed at higher levels in the brains of gamergates compared to workers (**Fig. 4D**, bottom). In fact, TFs in general displayed a gamergate-skewed expression profile: among 844 gamergate-biased genes, 79 (9.4%) were TFs, whereas only 27 out of 816 (3.3%) worker-biased genes were TFs (**Fig. 4E**). This three-fold over-representation of TFs among upregulated genes in the gamergate brains is consistent with a crucial role for transcriptional reprogramming during the behavioral transition (Gospocic et al., 2021).

Together, these observations suggest that behavioral reprogramming of workers into gamergates is supported by the upregulation of a disproportionate number of TFs, which may orchestrate the social transition by facilitating gamergate-specific chromatin loops.

### Gamergate-biased genes are controlled by super-interacting promoters

Given the increased expression of TFs in gamergate brains, we reasoned that some of them likely regulate caste-specific genes. In line with this, promoters activated in gamergate brains were strongly enriched for motifs of several gamergate-specific TFs (**Fig. 5A**), as well as for two worker-expressed transcriptional repressors (E(spl)mγ-HLH and dpn). The motif bound by lmd, a TF that regulates the insulin pathway (Park et al., 2014), which is crucial to caste differentiation, was enriched at gamergate promoters (**Fig. 5A**, **S7A**). We also noted that gamergate-specific promoters with Imd motifs formed multiple chromatin loops contacting enhancers (**Fig. 5B**). For example, the promoter of gamergate-specific gene *Oatp74D*, which regulates ecdysone transport across the blood-brain barrier in *Drosophila* (Okamoto and Yamanaka, 2020), connected to multiple intronic enhancers (**Fig. 5C**). Interestingly, the *Oatp74D* promoter formed 12 E–P loops in gamergate brains, and 8 (67%) of them were retained in worker brains although the gene was expressed at lower levels in workers (**Fig. 5C**).

**Figure 5.**
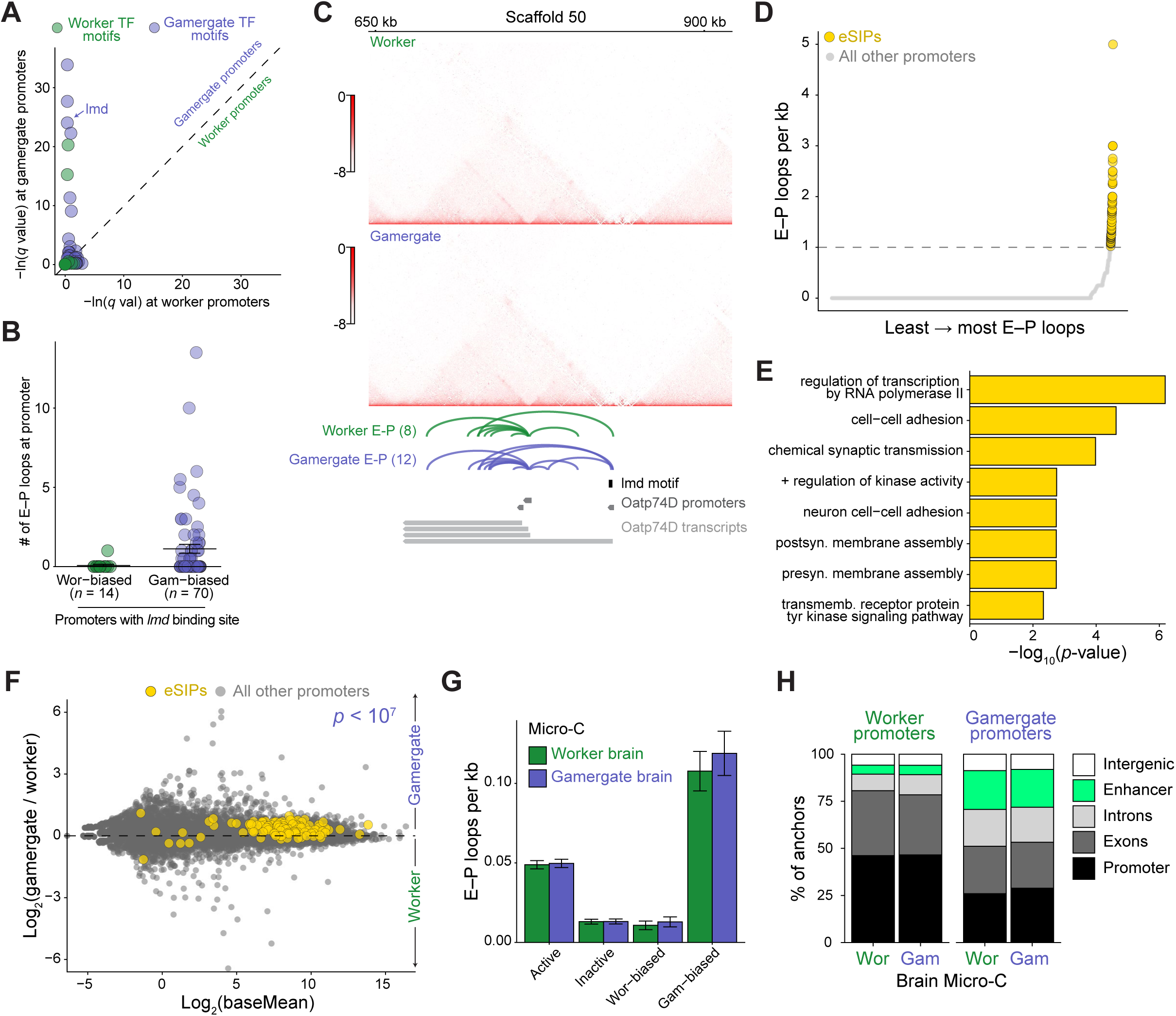
Gamergate-biased promoters are hyper-connected to enhancers. (A) Motif enrichment for worker-specific (green) or gamergate-specific (blue) TFs at promoters of caste-specific genes. Log-transformed *q* value of enrichment at gamergate promoters is plotted on the *y* axis and at worker promoters on the *x* axis. (B) Number of E–P loops originating from worker– or gamergate-biased promoters containing one or more lmd motif(s). (C) Example of a gamergate-specific gene *Oatp74D* showing balanced, KR normalized Micro-C interactions in worker and gamergate brains (top), all E–P loops with an anchor in *Oatp74D* promoters in either caste shown as arcs, location of *Oat74D* promoters, and *lmd* motif location. (D) Number of E–P loops (per kb) originating from all promoters ranked from least to most loops. eSIPs (yellow circles) were defined as promoters that have one or more E–P loops per kb. (E) GO terms enriched in genes with eSIPs as defined in (D). All genes were used as the background. (F) Differential genes expression in day-30 gamergate brains (*n* = 17) *vs.* day-30 worker brains (*n* = 11). The position of genes associated with eSIPs is shown by the yellow circles. *P* value is from a Fisher’s exact test for the proportion of eSIPs significantly enriched in each caste, compared to the distribution of all significantly differentially expressed genes. (G) E–P loop frequency at promoters of active (*n* = 8,623), inactive (*n* = 3,725), worker-biased (*n* = 816), and gamergate-biased (*n* = 844) genes in worker and gamergate brains. (H) Identity of regions looping to promoters of worker-biased genes (left panel) or gamergate-biased genes (right panel), in the brains of workers (left bars) or gamergates (right bars).

In the human brain cortex, certain developmentally regulated, lineage-specific promoters display enhanced chromatin interactions and have been termed “super-interactive promoters” (SIPs) (Heffel et al., 2024; Song et al., 2020). We hypothesized that caste-specific genes in *Harpegnathos* might share this property. By ranking promoters by their interactivity, we identified 297 genes under the control of SIPs, which we defined as promoters that formed two or more loops per kilobase (**Fig. S7B**). We further identified 118 genes associated with “enhancer SIPs (eSIPs)”, defined as promoters engaging in at least one loop per kilobase specifically with enhancers (**Fig. 5D**). Both SIPs and eSIPs were associated with active genes (**Fig. S7C**) and eSIP-associated genes were enriched for GO terms relevant to neuronal development and function, including “*chemical synaptic transmission*” and “*neuron cell-cell adhesion*” (**Fig. 5E**).

Projecting eSIPs-associated genes onto the differentially expressed genes between castes, we noticed that 87% were upregulated in the gamergate brain, with 22% of all eSIPs exhibiting significant (adjusted *p* < 0.05) gamergate-biased expression, and none demonstrating significant upregulation in worker brains (**Fig. 5F**). In fact, promoters of genes upregulated in gamergates were much more likely to form 3D loops with distal regions (**Fig. S7D**), particularly with enhancers (**Fig. 5G**), as compared to worker-biased promoters. Remarkably, and similar to *Oatp74D*, the increased connectivity of these loci was also observed in the worker brains (**Fig. 5G, S7D**), indicating that the loops are stable in both castes and suggesting that they might poise crucial gamergate genes for activation during behavioral reprogramming. Consistent with this, when all types of chromatin loops were analyzed, gamergate-specific promoters formed more frequent contacts with enhancers than worker-specific promoters (**Fig. 5H**).

Together, these observations point to an inherent property of genes upregulated during the reprogramming of workers into gamergates: these gamergate-biased genes are enriched for TFs and many of their promoters exhibit heightened connectivity to other regulatory elements, especially enhancers. Along with the observation that gamergate brains form overall more chromatin loops, these results support a central role for enhancer regulation and 3D genome organization in *Harpegnathos* brain plasticity during social and behavioral reprogramming.

## DISCUSSION

The reprogrammable social and behavioral states observed in adult *Harpegnathos* ants make this species an attractive model for uncovering the epigenetic bases of brain plasticity. We generated a high-resolution multi-dimensional epigenomic atlas of the *Harpegnathos* brain, integrating 3D genome organization, chromatin accessibility, histone modifications, and gene expression. Our study establishes a genome organization reference map for the *Harpegnathos* brain, detailing classes of TADs and chromatin loops. Importantly, caste comparisons revealed that gamergates upregulate genes enriched for TFs and controlled by hyper-connected promoters (SIPs). Our findings highlight the multi-layered regulatory complexity governing gene expression dynamics in the brain during behavioral reprogramming.

### A comprehensive epigenomic atlas of an ant brain

To our knowledge, the datasets generated for this study represent the most comprehensive epigenomic atlas in any social insect brain. In recent years, a growing number of high quality genome assemblies and gene annotation have become available in many insect species, including social insects (Gao et al., 2020; Lee et al., 2025; McKenzie and Kronauer, 2018; Shields et al., 2021; Wallberg et al., 2019). These resources have enabled comparative genomics and transcriptomic analyses, but a sophisticated dissection of gene expression dynamics and their epigenetic regulation requires detailed annotations of CREs, especially enhancers, which are still largely lacking in most insects other than *Drosophila*.

Efforts have been made to annotate enhancers in insect species based on sequence information and computational prediction models alone (Asma et al., 2024). In a few cases, enhancer annotations have been generated using experimental chromatin profiles and functional assays, including accessibility and histone marks in the honeybee *Apis mellifera* (Lowe et al., 2022), chromatin accessibility only in the pharaoh ant *Monomorium pharaonis* (Wang et al., 2020), chromatin accessibility combined with a functional assay, STARR-seq, in the fire ant *Solenopsis invicta* (Jones et al., 2025), and STARR-seq in the sweat bee *Lasioglossum albipes* (Jones et al., 2024). Here, we combined chromatin accessibility and multiple histone modification profiles along with ChromHMM analysis to annotate enhancers and their activation state in the *Harpegnathos* brain. Most importantly, and for the first time in a social insect, we utilized Micro-C to define their 3D connectivity.

While genome-wide chromatin conformation datasets have been reported for several insects, including *M. pharaonis* (Gao et al., 2020), *A. mellifera* (Jin et al., 2023), and multiple bumblebee species (Sun et al., 2021), only one was generated specifically from the brain, in *Drosophila* (Mohana et al., 2023), where long-distance meta-loops were first reported. To our knowledge, loops inferred from chromosome conformation data have not been used to assign target genes to enhancers on a genome-wide scale in any of these studies.

Performing Micro-C in *Harpegnathos* brains allowed us to both improve the genome assembly and link enhancers to distal promoters (**Fig. 1**). Almost half (797, 42%) of enhancers for which we could detect E–P loops (1,872) contacted one or more distal promoters. This is a relatively small fraction of all annotated enhancers (26,973), but it E–P interactions are notably difficult to capture (Lee et al., 2022) and our findings are consistent with reports in other species (Gasperini et al., 2019; Ing-Simmons et al., 2021; Lee et al., 2022). Nonetheless, examples of distal gene regulation are abundant in mammalian systems (Lettice et al., 2003; Stadhouders et al., 2012) and *Drosophila* (Ghavi-Helm et al., 2014), and knowledge of these E–P interactions in *Harpegnathos* brains will be crucial to correctly interpreting the gene regulatory networks that underpin the social transition and associated behavioral reprogramming.

### Characterization of TADs and chromatin loops in the *Harpegnathos* brain

Boundaries separating the 4,608 TADs in the *Harpegnathos* brain showed a substantial overlap with active promoters marked by accessible chromatin and H3K4me3 (**Fig. 2**). This feature is conserved across human (Dixon et al., 2012; Krietenstein et al., 2020), mice (Bonev et al., 2017; Dixon et al., 2012; Hsieh et al., 2020), flies (Hou et al., 2012; Ramirez et al., 2018; Sexton et al., 2012), yeast (Hsieh et al., 2015; Hsieh et al., 2016), and *Arabidopsis* (Sun et al., 2024; Wang et al., 2015). However, unlike in mammals, where TAD boundaries are predominantly associated with CTCF occupancy, *Harpegnathos* TAD boundaries showed no enrichment for CTCF motifs, similar to what has been observed in *Drosophila* (Acemel and Lupianez, 2023; Dixon et al., 2012; Ramirez et al., 2018). TAD boundaries in *Harpegnathos* were instead enriched for motifs recognized by other TFs, which varied depending on the boundary localization (i.e. genic vs. intergenic). Further classification of TADs according to their chromatin state revealed that the largest fraction of the genome is organized into “bivalent” TADs, which displayed both repressive and activating chromatin marks and were enriched for enhancers in various activation states (**Figure 3, S4**). This is notably different from *Drosophila* embryos and S2 cells, where the majority of the genome folded into inactive (or null/void) TADs (Ramirez et al., 2018; Sexton et al., 2012). While this and other observations in this study support an increased *cis*-regulatory complexity of the *Harpegnathos* genome, particularly within the brain, we cannot exclude that tissue of origin or other experimental factors might explain the observation.

Motif enrichment at loop anchors highlighted the importance of GAF, which functions as a pioneer-like TF and looping factor in *Drosophila* (Gaskill et al., 2021; Li et al., 2023), as well as additional TFs uniquely associated with *Harpegnathos* loop anchors. As an important computational control, we noted that BEAF-32, the most enriched TF motif at *Drosophila* anchors, was not present at *Harpegnathos* anchors. This is consistent with the absence of BEAF-32 homologs in the *Harpegnathos* genome and confirms the accuracy of our motif enrichment analyses.

### Increased 3D chromatin connectivity of gamergate brains and promoters

After 30 days of transition, gamergates selectively upregulated a disproportionate number of TFs in the brain compared to workers, many of which recognized motifs enriched at gamergate-specific 3D chromatin loops. Given the idea that TFs can act as anchors for chromatin loops to bring regulatory elements into proximity (Kim and Shendure, 2019), our finding suggests that some of these gamergate TFs are required to form new, gamergate-specific chromatin loops that underpin gene regulatory networks during behavioral reprogramming.

We identified a set of promoters with unusually high 3D connectivity (SIPs/eSIPs) (**Fig. 5**). These promoters were associated with genes encoding factors enriched for neuronal functions, suggesting that their tight regulation and fine-tuning in the brain relies on extensive proximal and distal *cis* regulation. Consistent with this, genes regulated during the social transition, in particular those activated by reprogramming of young workers into the distinct gamergate behavioral state, were strongly enriched for eSIPs.

SIPs were first defined in the human cortex, where they drive transcription of lineage-determining genes and TFs in a cell-type specific manner (Song et al., 2020). Our observations in *Harpegnathos* suggest that socially regulated genes in ants are finely regulated in a manner analogous to SIP-controlled genes in the human brain. Remarkably, these gamergate SIPs retain their hyper-connectivity even in the chromatin of worker brains, where those genes are poorly expressed. This argues that their 3D organization and dense regulatory network is not just a consequence of increased gene expression, but rather a reflection of the need for precisely tuned regulatory inputs at these loci during behavioral and social reprogramming.

Previous studies and our own anecdotal observations suggest that younger *Harpegnathos* workers are more responsive to caste transition (Opachaloemphan et al., 2021; Opachaloemphan et al., 2018), indicating that brain plasticity declines with age even in a highly reprogrammable species. We speculate that loss of the pre-established 3D architecture at gamergate-specific socially regulated promoters could contribute to the loss of behavioral plasticity observed in older ants, and possibly in other species.

## MATERIALS AND METHODS

### Harpegnathos saltator colony maintenance

*Harpegnathos saltator* colonies were housed in polystyrene boxes (Pioneer plastic) with a nest base made of dental plaster. Colonies were kept in a temperature-controlled (25°C) facility on a natural light/dark cycle. Ants were fed 2–3 times per week with live crickets (*Gryllodes sigillatus*). Colonies had light and dark chambers, wherein the nest with the brood was located in the dark chamber, while workers hunted crickets in the light chamber. Detailed procedures for *Harpegnathos* husbandry and colony maintenance are described in Moreno-Medina et al. (2026b).

### Worker to gamergate transition

We performed worker–gamergate transitions as previously described (Gospocic et al., 2021; Gospocic et al., 2017; Sheng et al., 2020), with minor modifications. In short, 20–25 callow females (4–13 days old; for each experiment ants were within a maximum of 3 day age difference) were isolated from their original colonies and transferred with 3–6 males to a new nest. Every female ant was painted with a unique two-color combination. The worker or gamergate status of transitioning ants was defined based on their behavior scores and ovary scores (see below). Detailed procedures for worker to gamergate transition are described in Moreno-Medina et al. (2026b).

### Behavior and ovary scoring

Ants were observed for three consecutive days, at least 1.5 hours per day, prior to collection on day 30. On each observation day, individual behavior scores were quantified. Gamergate-specific behaviors were assigned positive values: egg-laying (+3), dueling (+2). Worker-specific behaviors were assigned negative values: exiting nest (–0.5), defending (–0.5), nursing or cleaning (−2), and hunting (−2). Ants with conflicting behavioral observations (e.g. worker-like behavior on one day and or gamergate-like behavior on another day) were not further considered. For each individual, the raw scores were summed over the three observation days, resulting in a theoretical range from –15 to +15.

On collection day, ants were sacrificed, and their brains and ovaries were dissected. Their worker or gamergate status was confirmed by observing the ovaries. An ovary scoring system was established based on both the number and size of yolky oocytes present in all eight ovarioles. A yolky oocyte with a diameter > 250 µm but < 500 µm was assigned a score of +0.5, and a yolky oocyte with a diameter > 500 µm was assigned a score of +1, resulting in a raw score theoretically ranging from a minimum of 0 to a maximum of 8. Samples displaying discordant behavior and ovary phenotypes (e.g. worker-like behavior with enlarged ovary, or vice versa) were discarded. For plotting (**Fig. S6A**), both the raw behavior score and raw ovary score for each ant were linearly transformed to a standardized scale ranging from 0 to 10, using the formula 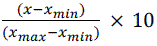. Detailed procedures for behavior scoring and ovary dissection are described in Moreno-Medina et al. (2026a).

### Isolation of nuclei from brains

Brains were dissected and washed in HBSS with 0.5% BSA (Sigma-Aldrich, A8577). Non-visual brains were obtained by removing optic lobes. This tissue-sampling allowed us to focus our analysis on regions of the brain more likely to encode higher functions such as learning, memory, and social behavior (Heisenberg, 2003). Detailed procedures for brain dissection are described in Moreno-Medina et al. (2026a).

First, freshly dissected brains were incubated for 10 min on ice in 120 µL nuclei isolating buffer containing 10 mM HEPES-NaOH pH 7.4, 10 mM KCl, 2.5 mM MgCl_2_, 0.1% Triton X-100, and 0.1% BSA. To disrupt the tissue and release nuclei, tissues were pipetted up and down through a 200 μL gel loading tip (Sarstedt, 70.1190.100) approximately 30 times. Samples were then passed through a 10 µm nuclei strainer (cellTrics, 04-004-2324) and nuclei were pelleted at 500 g for 5 min at 4°C. The pellet was resuspended in nuclei resuspension buffer containing 10 mM HEPES-NaOH pH 7.4, 90 mM KCl, 40 mM NaCl, 2.5mM MgCl_2_, 0.2 U/µL SUPERase Inhibitor (Thermo Fisher, AM2694), 1x protease inhibitor (Sigma-Aldrich, 11873580001), and 0.1% BSA. Nuclei were counted by Hoechst 33342 stain (Biotium, 40046) using a hemocytometer under a fluorescence microscope (ZOE Fluorescent Cell Imager, Bio-RAD). Detailed procedures for nuclei isolation are described in Kuang et al. (2026a).

### CUT&Tag

CUT&Tag was performed on freshly isolated nuclei following an adapted protocol based on Kaya-Okur et al. (2020); Kaya-Okur et al. (2019) and EpiCypher (2021). Each bulk reaction started with roughly 50,000 ant nuclei in 50 µL Wash150 buffer containing 20 mM HEPES-NaOH pH 7.5, 150 mM NaCl, 0.5 mM spermidine (Sigma-Aldrich, S2501-1G), and 1x EDTA-free Protease Inhibitor Cocktail (Sigma-Aldrich, 11873580001). 10 µL of Concanavalin A beads (Polysciences, 86057-3) in bead activation buffer (20 mM HEPES-NaOH pH 7.9, 10 mM KCl, 1 mM CaCl_2_, 1 mM MnCl_2_) was added to each sample and incubated on a nutator for 20 min at room temperature to allow binding to the nuclei. The nuclei bound to the beads were pelleted on a magnet and resuspended in antibody150 buffer (Wash150 supplemented with 0.1% BSA and 2 mM EDTA) containing the desired primary antibody at manufacturer recommended concentrations (see antibody list below). Samples were incubated on a nutator overnight at 4°C.

On the second day, the beads were pelleted on a magnet and resuspended in Wash150 supplemented with secondary antibody, and incubated for 60 min on a nutator at room temperature. After two washes in Wash150, samples were incubated with 0.034 µM homemade pG-Tn5 (purified according to Picelli et al. (2014)) in Wash300 buffer (20 mM HEPES-NaOH pH 7.5, 300 mM NaCl, 0.5 mM spermidine, 1x EDTA-free Protease Inhibitor Cocktail) for 60 min binding on a nutator at room temperature. After pG-Tn5 binding, three washes were performed to eliminate unbound pG-Tn5. Tagmentation was induced by exchanging with tagmentation buffer (Wash300 supplemented with 10 mM MgCl_2_) and samples were incubated for 60 min at 37°C. Nuclei were washed twice in 10 mM TAPS buffer (pH 8.5) with 0.2 mM EDTA. Samples were then resuspended in 5 µL SDS release buffer (10 mM TAPS, 0.1% SDS) and incubated overnight at 58°C to release the DNA fragments. For PCR, SDS quench buffer (0.67% Triton-X) was first added to neutralize SDS, and the nuclei-ConA bead lysate was used as input sample for direct amplification. The subsequent PCR reaction was performed using NEBNext HighFidelity 2x PCR Master Mix (New England Biolabs, M0541S) and Illumina Nextera i5 and i7 adapters to amplify the libraries for 14–16 cycles. The final libraries were purified using SPRI beads (1.3x), quantified by NEBNext® Library Quant Kit for Illumina (E7630) and sequenced on an illumina NovaSeq.

Antibodies used for bulk CUT&Tag were as listed and diluted according to manufacturer’s recommendations: H3K27me3 (rabbit, Epicypher, 13-0055), H3K27me3 rabbit, Cell Signaling, C36B11), H3K4me3 (rabbit, Epicypher, 13-0041), H3K4me3 (rabbit, Abcam, ab213224), H3K4me1 (rabbit, Abcam, ab8895), H3K27ac (rabbit, Abcam, ab4729), H3K9me3 (rabbit, Abcam, ab8898), H2AK119ub1, (rabbit, Cell Signaling, D27C4), H3K36me3, (rabbit, Abcam, Ab9050), Rabbit IgG (secondary antibody, guinea pig, antibodies-online, ABIN101961), IgG (negative control, rabbit, Epicypher, 13-0042). Detailed procedures for CUT&Tag using *Harpegnathos* brain nuclei are described in Kuang et al. (2026a).

### ATAC-seq

Bulk ATAC-seq was performed on freshly isolated nuclei following an adapted protocol based on Omni-ATAC (Corces et al., 2017; Grandi et al., 2022). Roughly 30,000–50,000 *Harpegnathos* brain nuclei were used for each ATAC-seq reaction by first pelleting the nuclei and resuspending in 50 µL ATAC master mix (10 mM Tris-HCl pH 7.5, 5 mM MgCl_2_, 10% dimethylformamide, 0.01% digitonin, 0.1% Tween 20, 0.2 µM homemade Tn5 purified according to Picelli et al. (2014)). Samples were incubated for 30 min at 37°C in a thermomixer at 1,000 rpm, and then cleaned up using ChIP DNA Clean & Concentrator to extract DNA fragments. The subsequent PCR reaction was performed using KAPA HiFi HotStart ReadyMix (2X) (Roche, KK2601) and Illumina Nextera i5 and i7 adapters to amplify the libraries for 18 cycles. The final libraries were purified using SPRI beads (1.3x) (Beckman Coulter, B23318), quantified by NEBNext® Library Quant Kit for Illumina (E7630) and sequenced on an Illumina NextSeq.

### Brain RNA-seq

For each biological replicate, a single non-visual brain was used. Freshly dissected *Harpegnathos* non-visual brains were lysed and homogenized in TRIzol for RNA-purification. For library preparation, polyA+ RNA was enriched using Dynabeads Oligo(dT)25 (ThermoFisher, 65001). Strand-specific libraries were prepared using NEBNext® Ultra™ II Directional RNA Library Prep Kit (NEB, E7645S) and the steps included RNA fragmentation and random priming, first strand cDNA synthesis, second strand cDNA synthesis, end repair and 5’ phosphorylation, U excision, and PCR enrichment. The final libraries were purified using SPRI beads (1x) (Beckman Coulter, B23318), quantified by NEBNext® Library Quant Kit for Illumina (E7630).

### Micro-C

Micro-C was performed based on an optimized protocol from Slobodyanyuk et al. (2022). For each biological replicate, a single non-visual brain was used. Freshly dissected *Harpegnathos* non-visual brains were fixed in PBST (0.1% Triton-X in PBS) containing 1% fresh formaldehyde (Thermo Fisher, 28906) for 15 min at room temperature. Fixation was quenched in 540 mM Tris-HCl pH 7.5 for 5 min and brains were washed 3 times in PBST. Brains were fixed further in 3 mM DSG (Thermo Fisher, A35392) diluted in PBST for 45 min at room temperature. Fixation was quenched in 540 mM Tris-HCl pH 7.5 for 5 min and brains were washed 3 times in PBST. Tissues were resuspended in MB#1 buffer (50 mM NaCl, 10 mM Tris, 5 mM MgCl_2_, 1 mM CaCl_2_, 0.2% NP-40, 1x protease inhibitor cocktail) and incubated on ice for 20 min. For homogenization, tissues were passed through a 200 μL gel loading tip by pipetting up and down about 30 rounds. The tissue homogenate was digested by 10 U of MNase in 100 µL of MB#1 buffer at 37°C, and digestion was stopped by incubating in 4 mM EGTA for 10 min at 65°C. After two washes in MB#2 buffer (50 mM NaCl, 10 mM Tris-HCl pH 7.5, 10 mM MgCl_2_), the pellet was resuspended in end-chewing buffer (1x MB#2, 4 mM ATP, 6 mM DTT) and incubated for 15 min at 37°C on a shaking thermomixer (1000 rpm for an interval of 15 s every 3 min). 5 μL of 5 U/μL Klenow fragment was added to each sample and incubated for 15 min at 37°C on a shaking thermomixer. 25 μL of “End-Labelling Mix” containing 1x T4 DNA ligase buffer (Enzymatics, L6030-HC-L), 0.2 mM biotin-14-dATP (Jena Bioscience, NU-835-BIO14-S), 0.2 mM biotin-14-dCTP (Jena Bioscience, NU-809-BIOX-S), 0.2 mM dTTP (Jena Bioscience, NU-1004L), 0.2 mM dGTP (Jena Bioscience, NU-1003L), and 0.1% BSA was then added to the sample for an incubation of 45 min at 25°C on a shaking Thermomixer. To inactivate the enzymes, EDTA was added to samples (final 30 mM). After washing in MB#3 buffer (500 mM Tris-HCl (pH 7.5), 100 mM MgCl₂, 1% BSA), samples were resuspended in 250 µL proximity-ligation mix containing 1x T4 DNA ligase buffer (Enzymatics, L6030-HC-L), T4 DNA Ligase (5000 U in 250 μL reaction) (Enzymatics, L6030-HC-L), and 0.1 mg/mL BSA and incubated at RT for 2.5 hours. To remove unligated ends, samples were pelleted again and resuspended in 100 µL exonuclease mix containing 1x NEBuffer 1 (NEB, B7001S) and Exonuclease III (500 U in 100 μL reaction) (NEB, M0206S) and incubated for 15 min at 37°C on a shaking thermomixer. Finally, for reverse-crosslinking, samples were pelleted and resuspended in 150 µL SDS-Proteinase K release mix containing 1% SDS, 0.5 mg/mL proteinase K (Thermo Fisher, AM2548), 0.1 mg/mL RNase A (Fisher Scientific, 10753721) and incubated overnight at 65°C.

DNA was released by adding 200 µL phenol:chloroform:isoamyl alcohol (PCIA) and cleaned up with a ZymoClean spin column (Zymo Research, D4007). Biotin-labeled DNA was incubated with and bound to Dynabeads MyOne Streptavidin C1 (Thermo Fisher, 65001) in 1x B&W buffer (5 mM Tris-HCl, 1 M NaCl, 0.5 mM EDTA) for 20 min at room temperature. Samples were then washed twice in 1x TBW buffer for 5 min at 55°C and rinsed once in 500 µL 10 mM Tris pH 7.5. For end repair and adapter ligation, DNA was first resuspended in a master mix prepared using NEBNext Ultra II DNA Library Prep Kit (3.5 µL reaction buffer, 1.5 µL enzyme mix, 25 µL H_2_O) (NEB, E7645S), and ligated to NEB Illumina universal adapters using components from NEBNext Ultra II Ligation Module (15 µL ligation master mix, 0.5 µL ligation enhancer) (NEB, E7595S). Afterwards, 1.5 μL NEB USER enzyme (NEB, M5505S) was added and samples were incubated for 15 min at 37°C. The reaction was washed in 1x B&W buffer once at 55°C for 5 min, rinsed once in 10 mM Tris pH 7.5, and beads were resuspended in 22 µL 10 mM Tris pH 7.5. For library amplification, the PCR reaction was assembled by adding 25 µL KAPA HiFi Hot Start Mix 2x (Roche, KK2601) and i5&i7 primers (300 nM) and amplified for 18 cycles. The library was finally purified with 1x SPRI beads, quantified by NEBNext® Library Quant Kit for Illumina (E7630), and sequenced on an illumina NextSeq. Detailed procedures for *Harpegnathos* brain Micro-C are described in Kuang et al. (2026b).

## DATA PROCESSING AND ANALYSIS

### Genome assembly and annotation

The *Hsal*_v9.0 genome was generated as described below and used for all analyses. Gene annotations from the current *Hsal*_v8.6 genome build (NCBI *Hsal*_v8.6 annotation, GCA_003227716.2) combined with Iso-Seq annotations produced in Shields et al. (2021), were lifted over to the *Hsal*_v9.0 genome, as detailed below. TFs were identified as any protein containing a known DNA-binding domain. Promoters were annotated as the regions ±1kb from transcription start sites, with overlapping promoters from the same gene merged.

GO analysis was performed using an in-house script comparing terms associated with genes in the test set with terms associated with the background, non-test set. *P* values were calculated from Fisher’s exact tests on this contingency matrix and adjusted using the Benjamini-Hochberg false discovery rate method. GO terms associated with each gene were obtained from the NCBI *gene2go* file, with descriptions from the Gene Ontology knowledgebase (Gene Ontology et al., 2023).

### Micro-C analysis

#### Update of genome assembly with Micro-C

164 million Micro-C reads from workers were processed using Juicer (Durand et al., 2016b) with no restriction site specified, mapping to the *Hsal*_v8.6 assembly. The resulting merged, deduplicated bam file *merged_dedup.bam* was converted to a *pre* format file using the sam_to_pre.awk utility. This input was used to run the 3D-DNA pipeline (Dudchenko et al., 2017) (https://github.com/aidenlab/3d-dna) with the parameter –r 0, which specifies 0 rounds of mismatch correction. The new assembly was called *Hsal*_v9.0. Gene annotations were lifted over from *Hsal*_v8.6 using the liftOver tool (Hinrichs et al., 2006) following creation of a chain file using axtChain, chainMergeSort, chainSplit, chainSort, chainNet, and netChainSubset.

#### Mapping, contact map generation, loop detection/differential loops

Micro-C reads obtained from 12 workers profiles (separate from the Micro-C samples used for genome assembly) and 13 gamergates profiles were mapped to the *Hsal*_v9.0 assembly with bwa mem (Li and Durbin, 2010). Pairtools (Open2C et al., 2023) parse, sort and dedup, and split were used to further process the files, with the parameters specified in the Dovetail Linked-Read Analysis Page (https://dovetail-analysis.readthedocs.io/en/latest/index.html). Juicer pre (Durand et al., 2016b) with KR and SCALE normalizations was used to convert the split pairs files into *.hic* files, which were converted to *cool* contact maps using hicConvertFormat (Ramirez et al., 2018; Wolff et al., 2018; Wolff et al., 2020) at resolutions of 200 bp, 400 bp, 800 bp, 1 kb, 5 kb, and 10 kb. These *cool* files were balanced with cooler balance (Abdennur and Mirny, 2020). Merged files were generated by merging the pairs files from replicates within castes with pairtools merge, then following the rest of the analysis with the merged pair files.

Loops were called for all resolutions with mustache (Roayaei Ardakany et al., 2020) with *p* value threshold 0.1. Loops from 200 bp, 400 bp, 800 bp, and 1 kb resolution were merged, keeping the loop with the highest (i.e. lowest bp) resolution in cases where a loop was called in multiple resolution. The resulting merged loops were categorized as P–P, P–E, E–E, P–GB, E–GB, GB–GB, or other (all other combinations of these categories, as well as loops with one anchor to a non-enhancer intergenic region). Differential loops between castes were called using diff_mustache.

#### Insulation score calculation, TADs

Insulation scores were calculated at resolutions from 200 bp to 10 kb with cooltools insulation (Open2C et al., 2024). Selecting 400 bp resolution as a balance between sensitivity and smoothing, TADs were identified as boundary regions with a boundary strength ≥ 1.

For A/B compartment assignment (**Fig. S3C**), eigen value decomposition was first performed using cooltools eigs-cis (Open2C et al., 2024) at 400 bp resolution. A/B compartments on the longest 30 scaffolds were assigned using manual curation of the compartments differentiated by the first eigenvalue. Regions with high H3K27ac and low H3K27me3 were assigned to the A compartment, while regions with high H3K27me3 and low H3K27ac were assigned to the B compartment.

#### Enhancer–promoter assignment

Enhancers were annotated using H3K4me1, H3K27ac, ATAC-seq, and H3K27me3 signal as described in the “enhancer annotation” section. The merged loops at resolution 200 bp, 400 bp, 800 bp, and 1 kb were used for this analysis, as anchors using 5 kb and 10 kb resolution often overlapping many elements, making it difficult to annotate specific features to each side of the loop. Anchors were allowed to overlap multiple enhancers or multiple promoters to be considered part of an enhancer-promoter loop. Enhancers with a loop anchor, with at least one promoter overlapping the other loop anchor, were assigned to those promoter(s).

#### Drosophila Micro-C and enhancer analysis

For comparison with *Drosophila*, much of the Micro-C analysis above was repeated using data from Mohana et al. (2023). All analyses, including read processing and mapping, contact map generation at various resolutions, TAD detection, loop detection and merging, and motif finding at boundary regions were performed the same way, mapping to dm6 instead of *Hsal*_v8.6. Enhancer annotations from REDfly (Keranen et al., 2022) were used for categorizing *Drosophila* loops.

#### Meta-loop analysis

Meta-loops in the *Harpegnathos* brain were identified using custom code provided in Mohana et al. (2023), specifically the meta_loops.R script. Default parameters were used, with the exceptions that only the largest 25 scaffolds were searched for meta-loops and the resolution was set to 5kb. From the 99 meta-loops identified from worker Micro-C, one was a intra-TAD loop and was discarded for other analysis.

### CUT&Tag and ATAC analysis

#### Mapping, peak calling, and IgG normalization

Adapters and low quality bases were trimmed from CUT&Tag and ATAC fastqs using CutAdapt (Martin, 2011) and TrimGalore (Krueger, 2024). The resulting reads were mapped to the *Hsal*_v9.0 genome with bowtie2, version 2.5.1 (Langmead and Salzberg, 2012). Duplicates were marked using Picard MarkDuplicates (Institute, 2019) and removed with samtools (Danecek et al., 2021). To filter out sub-nucleosomal peaks, reads with a fragment length less than 120bp were removed from all non-ATAC samples. Bigwig files were generated using DeepTools (Ramirez et al., 2016) with CPM normalization. Peaks were called with macs3 (Zhang et al., 2008), with narrow peaks called for ATAC, H3K27ac, and H3K4me3, and broad peaks called for H3K27me3, H3K4me1, H3K36me3, H2AK119ub, and H3K9me3.

IgG/ATAC contamination was present in different amounts for each antibody. An estimate of the amount of contamination was determined by first finding peaks present in the IgG samples but not in any CUT&Tag sample, reasoning that reads in these peaks would contain largely reads independent of antibody capture. About 0.27% of IgG reads were in these peaks. The percent of reads in these peaks was calculated for each IP sample, ranging from an average 0.008% of H3K27me3 reads to 0.42% of H3K9me3 reads. Scaling factors for IgG correction were calculated from comparing these percents to the IgG percents; samples with a higher percentage of expected contaminated reads would have more IgG subtraction during background correction. Scaled IgG corrected bigwigs were produced using these factors.

### ChromHMM

ChromHMM (Ernst and Kellis, 2012) was run for worker and gamergate using CUT&Tag and ATAC bams as input, with a merged IgG input from both castes used as background to minimize caste-specific differences driven by IgG, which tracked closely with ATAC. A bin size of 200 bp was used for BinarizeBam and 15 states were used in LearnModel. States were labeled manually using the emission parameters of each sample.

### Enhancer annotation

Consensus peaks were calculated for H3K27ac, ATAC, and H3K27me3 in workers, as well as H3K4me1 in both workers and gamergates, requiring the peak to appear in at least two replicates. Regions with consensus H3K4me1 peaks in either caste were classified using worker chromatin profiles into active (overlap with ATAC and/or H3K27ac peak, no overlap with H3K27me3 peaks), bivalent (overlap with ATAC and/or H3K27ac peak, and H3K27me3 peak), primed (no overlap with ATAC, H3K27ac, or H3K27me3 peaks), or poised (no overlap with ATAC or H3K27ac peaks, overlap with H3K27me3 peak). Enhancers were categorized as promoter-associated, exonic, intronic, or intergenic. Only intronic and intergenic peaks were retained as putative enhancers. Enhancers within 10 kb of a TSS were assigned to the nearest TSS. As described above, Micro-C from both castes was then used to update this initial proximity-based annotation for enhancer gene targets; if Micro-C linked an enhancer to a promoter that was not assigned by proximity, we gave priority to the 3D-connected promoter.

Additionally, a blacklist was constructed and any enhancers overlapping with the blacklist were removed for further analyses. Read counts on each 1 kb bin in the genome were produced using DeepTools multiBamSummary (Ramirez et al., 2016). For each replicate of CUT&Tag, including the IgG, and ATAC, the top decile of bins by counts were noted. Bins overlapping promoters were excluded from the analysis. Within each IP/ATAC, bins present in ≥ 90% of the replicates were considered top decile bins for that IP. Bins appearing in the top decile of IgG and at least 4 of the 9 samples (ATAC, H3K27ac, H3K4me3, H3K4me1, H3K36me3, H3K27me3, H2AK119ub, H3K9me3) were deemed regions of commonly high signal and formed the blacklist.

### RNA-seq analysis

As described above for the CUT&Tag/ATAC mapping, RNA-seq fastqs were trimmed with CutAdapt and TrimGalore. Resulting reads were mapped to the *Hsal*_v9.0 genome using STAR version 2.7.10a with parameters *--alignIntronMax 50000 –-outFilterScoreMinOverLread 0.9 –-outFilterMatchNminOverLread 0.9 –-outFilterMismatchNoverLmax 0.05 –-outFilterMatchNmin 16*. Duplicates were marked using Picard MarkDuplicates (2019). Counts on genes were calculated with an in-house R script utilizing the GenomicRanges (ver 1.50.2) (Lawrence et al., 2013) functions *assay* and *summarizeOverlaps*, ignoring secondary alignments and running in paired-end mode. Differential expression was performed with DESeq2 (Love et al., 2014).

### Motif finding

TAD boundaries were separated into promoter, genic, and intergenic based on overlaps relevant genomic features. Sequences were tested for enrichment over shuffled input sequences, preserving dinucleotide frequencies, with the MEME suite tool Simple Enrichment Analysis, or SEA (Bailey et al., 2015).

Loop anchors were tested against shuffled genome regions, promoters with loops were tested against promoters without loops, while promoters with loops and a H3K4me3 peak were tested against promoters with loops without a H3K4me3 peak, all using SEA as above, to find motif enrichment of these genomic features.

Motif enrichment for *Drosophila* promoters with a loop (using Micro-C data from Mohana et al. (2023)) and H3K4me3 peak (using data from Rickels et al. (2017)) was performed with SEA as above, comparing to promoters with a loop but lacking a H3K4me3 peak. All motifs with a *q* value < 0.05 were considered enriched.

For detection of promoters with the *lmd* motif, MEME suite tool FIMO (Bailey et al., 2015) was run on all promoters. All occurrences with a *p* value < 10^-5^ were considered significant.

### Visualization

Genome-wide and finer resolution Micro-C contact maps were visualized using Juicebox (Durand et al., 2016a). Micro-C contact maps, insulation scores, CUT&Tag tracks, RNA-seq tracks, TADs annotations, and genomic features were visualized with CoolBox (Xu et al., 2021). Loops connecting scaffolds or genomic features were visualized using IGV (Robinson et al., 2023).

All plots were generated with ggplot2 (ver 3.4.2) (Wickham, 2016) and heatmaps were created using pheatmap (ver 1.0.12) (Kolde, 2025). The viridis color palette (ver 0.6.2) (Garnier et al., 2024) was used for some visualizations.

## DATA AVAILABILITY

Sequencing data were deposited in GEO with the accession number GSE325400. The new Hsal 9.0 assembly is available on GenBank with accession number JBWJFF000000000.

## ACKNOWLEDGMENTS

R.B. acknowledges the funding from Max Planck-Humboldt Research Award 2020, lab space at Uniklinik Freiburg, and resources shared with the Timmers lab. M.K. acknowledges the support from the International Max Planck Research School for Epigenetics, Biophysics and Metabolism.

## AUTHOR CONTRIBUTIONS

Conceptualization: R.B.; data collection: M.K., A.A.; methodology: M.K., A.A., S.M.M., E.J.S., R.B.; reagent generation: J.D., K.S.; investigation: M.K., E.J.S., R.B.; formal analyses: E.J.S.; writing – original draft: M.K., E.J.S., and R.B.; writing – review & editing: all authors; visualization: M.K., E.J.S., R.B.; supervision: R.B.; funding acquisition: R.B., H.T.M.T, M.P.

## SUPPLEMENTARY FIGURE LEGENDS

**Figure S1.**
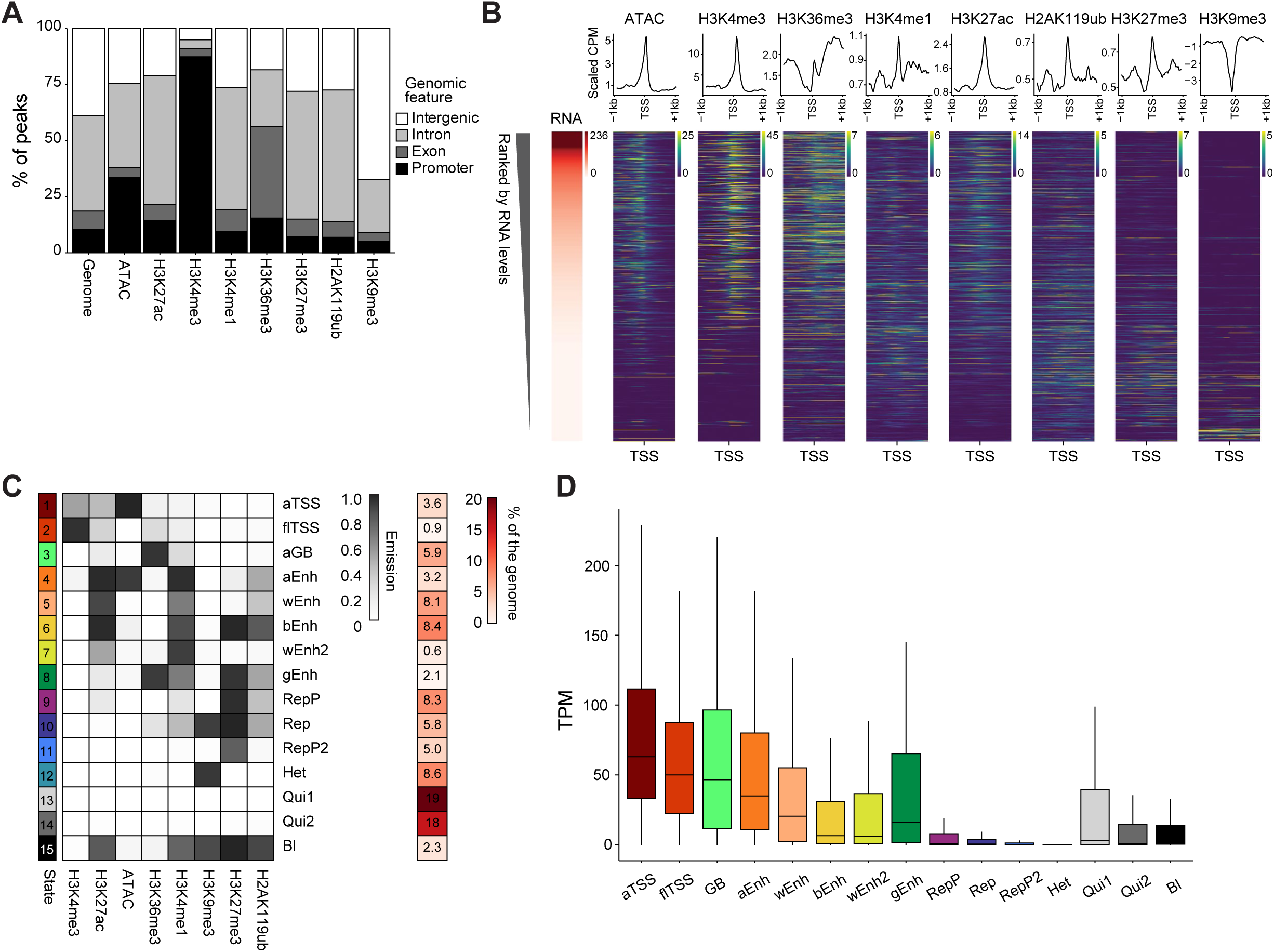
Chromatin accessibility and histone modifications, related to Figure 1. (A) Relative distribution of consensus chromatin peaks (in %) among the indicated genomic features. Promoters were defined as TSS ± 1 kb. Feature coverage on the whole genome is shown as background control. Consensus peaks for each IP were defined as regions where at least two biological replicates for each mark displayed a peak. (B) ATAC-seq (CPMs) or CUT&Tag for the indicated histone modifications (scaled IgG-normalized signal) at the TSS ± 1 kb for 17,633 genes. The heatmaps were all sorted according to RNA levels (TPMs) shown on the left. (C) Chromatin states defined by ChromHMM. The emission heatmap on the left shows the frequency of histone modifications and chromatin accessibility in each of the 15 chromatin states, whereas the portion of the genome assigned to each state is shown on the right (in %). States were annotated based on histone modification patterns: aTSS, active TSS; flTSS, flanking TSS; aGB, active gene body; aEnh, active enhancer; wEnh, weak enhancer; bEnh, bivalent enhancer; gEnh, genic enhancer; RepP, Polycomb-repressed; Rep, repressed; Het, constitutive heterochromatin; Qui, quiescent; Bl, blacklist. (D) Expression in TPM of genes whose promoters belong to each chromatin state defined in (C).

**Figure S2.**
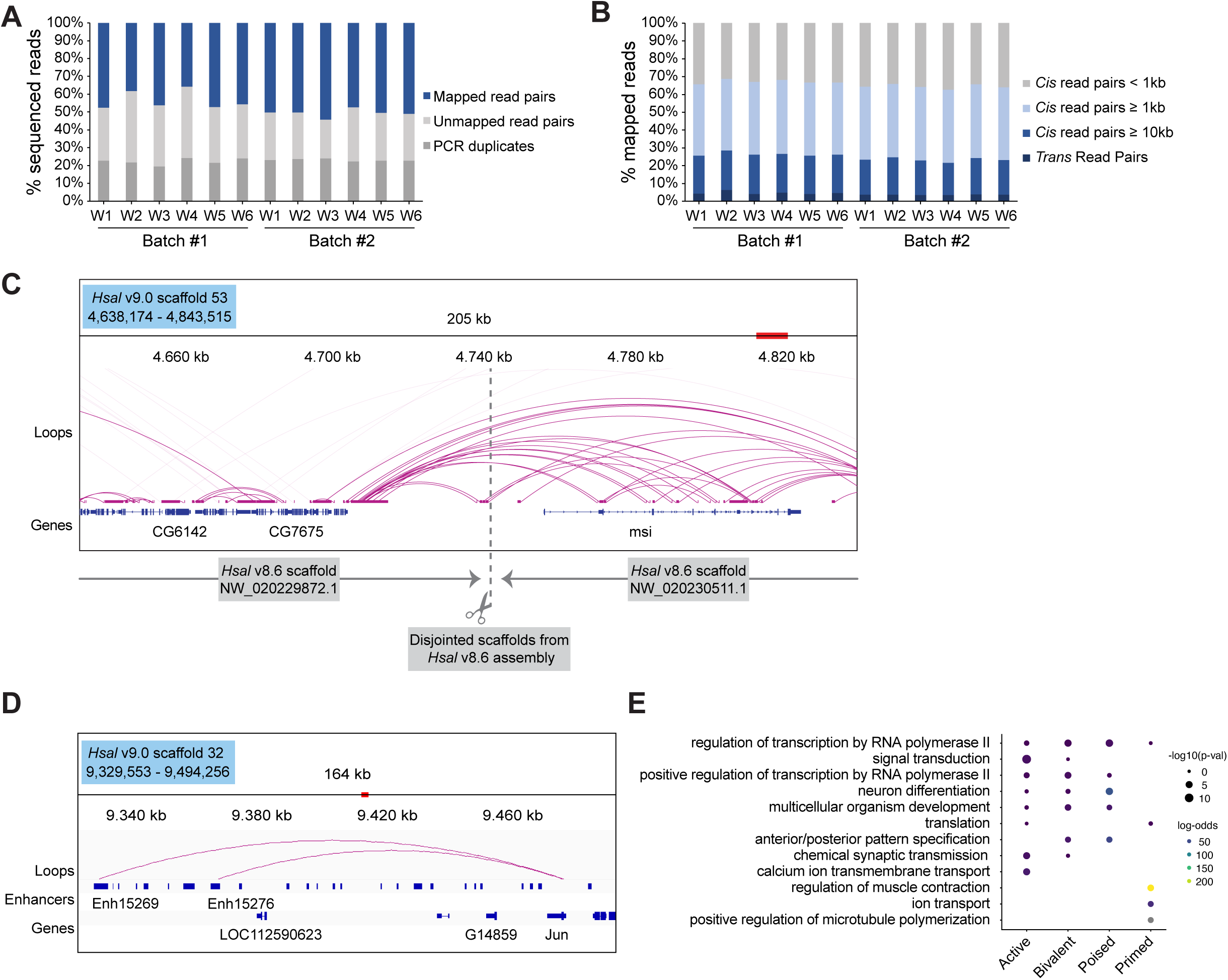
Quality assessment of Micro-C and example loci, related to Figure 1. (A) Quality metrics for Micro-C mapping. Mapped read pairs (blue), unmapped read pairs (light gray), and PCR duplicated read pairs (dark gray) as a % of total reads are shown for the 12 biological replicates, obtained in two batches. W1, worker sample 1, etc. (B) Types of interactions obtained from Micro-C sequencing, classified into *trans* read pairs (interaction between different scaffolds) or *cis* read pairs with interaction distances of < 1 kb (gray), between 1 and 10 kb (light blue), or larger than 10 kb (dark blue). (C) Two scaffolds from the *Hsal* v8.6 assembly were joined in *Hsal* 9.0 based on connecting Micro-C read pairs, shown as loops. (D) Example of E–P loops connecting the *Jun* promoter to two distal enhancers, which are separated from the target promoter by multiple intervening genes and > 100 kb in linear sequence. (E) GO enrichment analysis for genes assigned by Micro-C to active (*n* = 7,184), bivalent (*n* = 8,672), poised (*n* = 4,999), and primed enhancers (*n* = 6,737) in worker brains.

**Figure S3.**
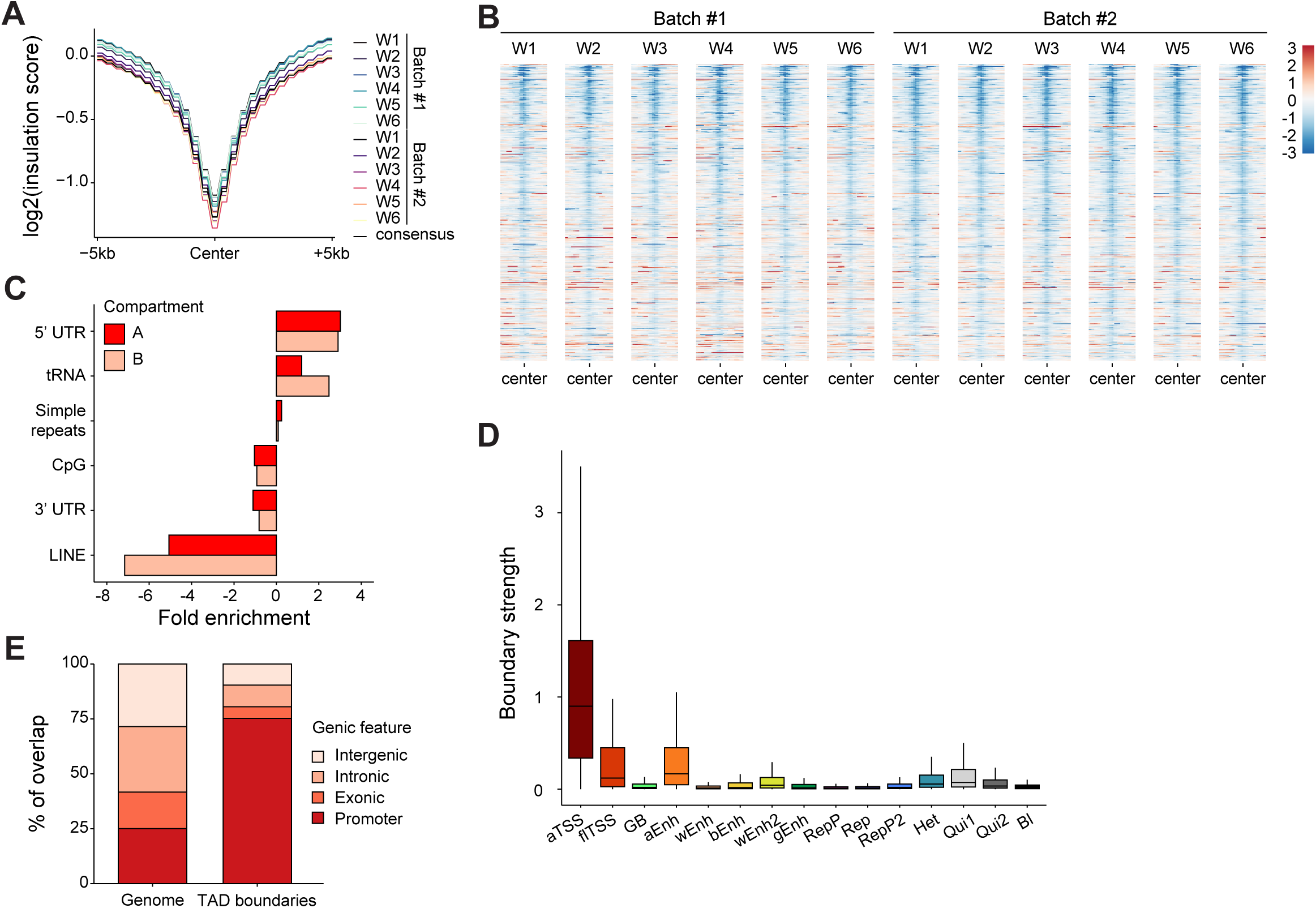
Additional characterization of TAD boundaries, related to Figure 2. (A) Insulation scores centered at TAD boundaries. (B) Insulation scores at all TAD boundaries, sorted by boundary strength from consensus TAD calling for all Micro-C biological replicates. Heatmaps are centered at TAD boundaries. (C) Enrichment of TAD boundaries overlapping genic features compared to the background genome, divided by the total width of each genic feature in the genome. (D) Boundary strength for TADs grouped by ChromHMM states. (E) Percentage of *Drosophila* TAD boundaries overlapping with promoter (± 1 kb from the TSS), exonic, intronic, and intergenic regions, compared to the composition of the background genome. Analysis performed on published *Drosophila* brain Micro-C data from Mohana et al., 2023.

**Figure S4.**
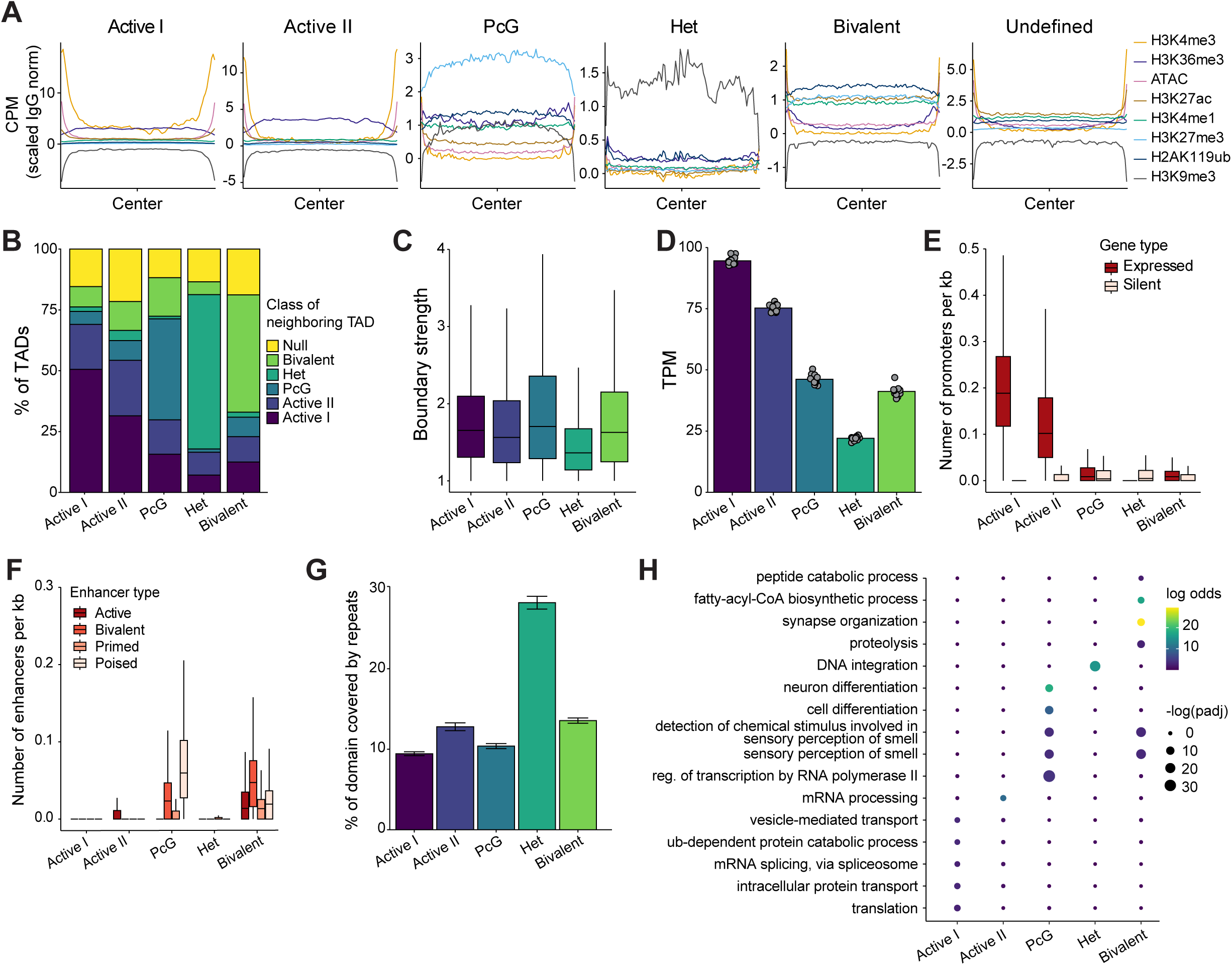
Additional analyses on TAD classes, related to Figure 3. (A) Metaplots showing ATAC-seq (CPM) and CUT&Tag signal (H3K4me3, H3K27ac, H3K4me1, H3K27me3, H2AK119ub, and H3K9me3, normalized with scaled IgG) across TADs classified according to their chromatin state. (B) For each TAD class defined in Fig. 3A, percentage of neighboring TADs from each class. (C) Boundary strengths for TADs classified according to their chromatin state. (D) Expression in TPM of genes within each TAD class. Grey dots show 13 RNA-seq biological replicates. (E) Promoter density per kb in each TAD class. Promoters are separated into expressed (TPM ≥ 10, red) and silent (TPM ≤ 1, pink). (F) Enhancer density per kb in each TAD class. Enhancers are classified according to their chromatin state, as defined in Fig. 1E. (G) Percentage of TADs from each class occupied by major repeat classes (LINEs, LTRs, LCRs, STRs, and DNA repeats). (H) GO terms enriched in genes within each TAD class.

**Figure S5.**
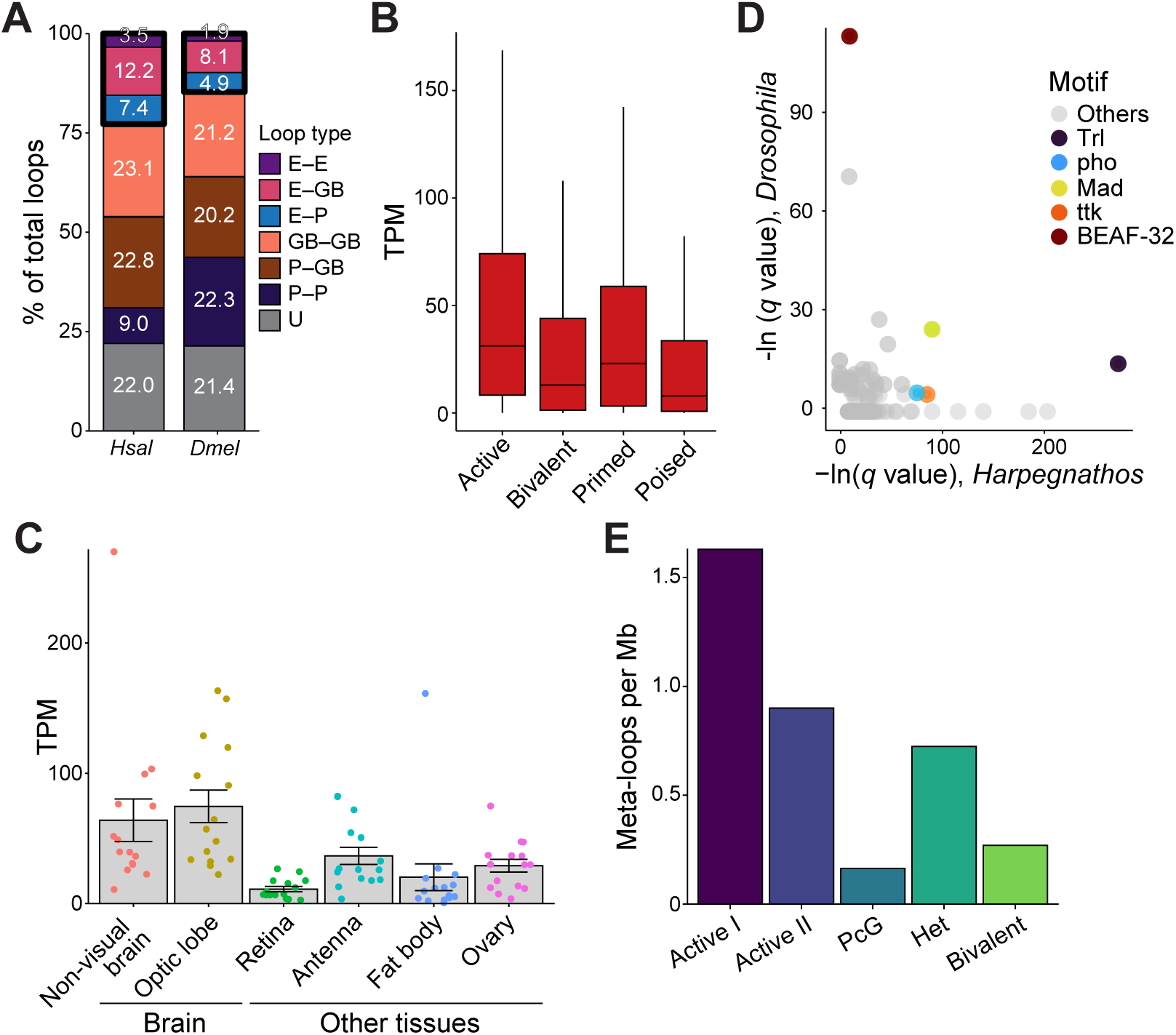
Additional analyses on loops and meta-loops, related to Figure 3. (A) Percentage of total loops categorized as enhancer–enhancer (E–E), enhancer–gene body (E–GB), enhancer–promoter (E–P), gene body–gene body (GB–GB), promoter–gene body (P–GB), promoter–promoter (P–P), or undefined (U), in *Harpegnathos* and *Drosophila*. *Drosophila* Micro-C data from adult brains (Mohana et al., 2023) was reanalyzed with enhancers defined by REDfly (Keranen et al., 2022). The thick black outline highlights loops with at least one anchor at enhancers. (B) Expression in TPM of genes contacting active, bivalent, primed, and poised enhancers. (C) Expression of TFs in various *Harpegnathos* tissues. Each dot represents a *Harpegnathos* homolog for one of the TFs shown in Fig. 3F. RNA-seq data from Shields et al. (2018). (D) Motif enrichment at looped promoters with H3K4me3 against looped promoters without H3K4me3, represented as –ln(*q* value), in *Harpegnathos* (*x* axis, same as Fig. 3F, right column) vs. *Drosophila* (*y* axis). Select motifs enriched in *Harpegnathos* are highlighted, along with BEAF-32. (E) Meta-loops per Mb in each TAD class.

**Figure S6.**
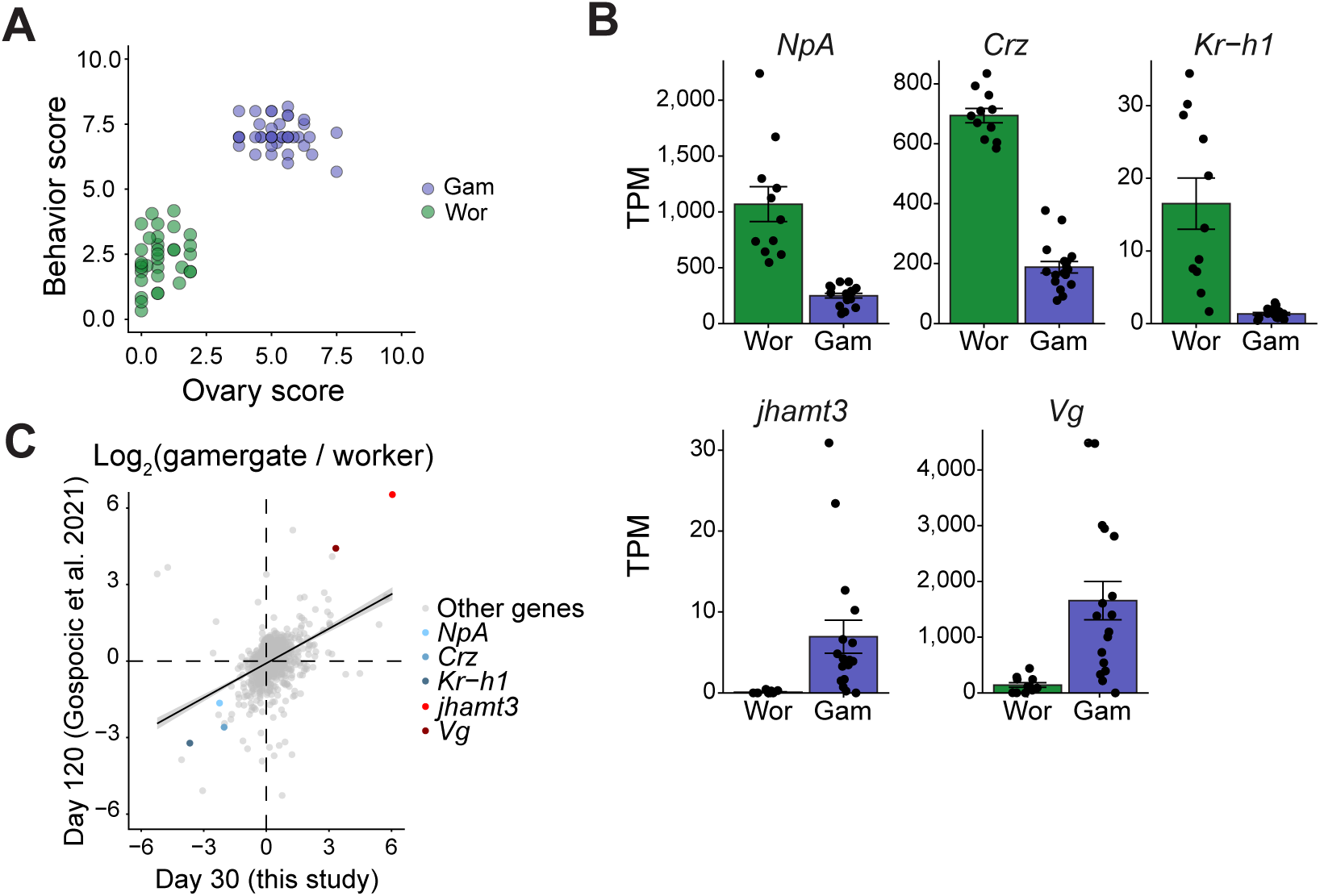
RNA-seq in worker and gamergate brains, related to Figure 4. (A) Correlation between behavior scores and ovary scores for workers (*n* = 34) and gamergates (*n* = 44) used in this study. (B) RNA levels (TPM) of genes identified in previous studies as worker-biased (top) or gamergate-biased (bottom) in worker (*n* = 11) and gamergate (*n* = 17) brains. (C) Correlation between log-fold-changes between RNAs in gamergate and worker brains, comparing day-30 transitions (this study) with day-120 transitions (Gospocic et al., 2021). All genes differentially expressed (adjusted *p* < 0.05) in either dataset are shown, with caste-biased genes from (B) highlighted.

**Figure S7.**
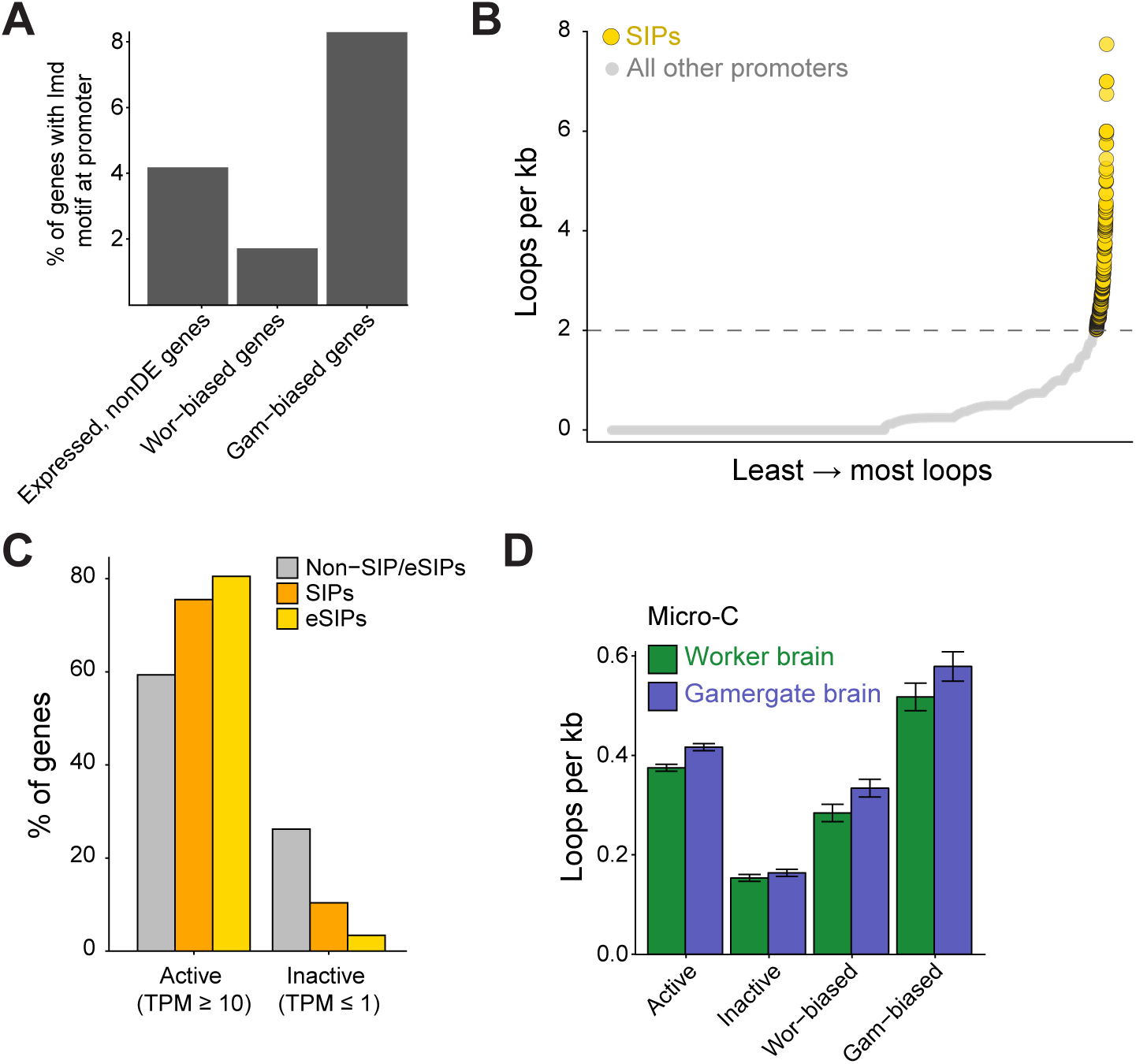
Additional analyses on SIPs and eSIPs, related to Figure 5. (A) Percentage of expressed, nonDE (*n* = 7,153), worker-biased (*n* = 816), and gamergate-biased genes (*n* = 844) with one or more occurrence(s) of the lmd motif at their promoter. (B) Number of loops (per kb) originating from all promoters ranked from least to most loops. SIPs (yellow circles) were defined as promoters that have two or more loops per kb. (C) Percentage of eSIPs, SIPs, and non-SIP/non-eSIP genes considered active (TPM ≥ 10) or inactive (TPM ≤ 1). (D) Looping frequency of all loops at promoters of different gene categories in worker and gamergate brains.

## SUPPLEMENTARY TABLE LEGENDS

**Table S1.** Micro-C sequencing reads and quality metrics for worker samples.

**Table S2.** Micro-C sequencing reads and quality metrics for gamergate samples.

## REFERENCES

1. Abdennur, N., and Mirny, L.A. (2020). Cooler: scalable storage for Hi-C data and other genomically labeled arrays. Bioinformatics 36, 311–316.

2. Acemel, R.D., and Lupianez, D.G. (2023). Evolution of 3D chromatin organization at different scales. Curr Opin Genet Dev 78, 102019.

3. Alberini, C.M. (2009). Transcription factors in long-term memory and synaptic plasticity. Physiol Rev 89, 121–145.

4. Asma, H., Tieke, E., Deem, K.D., Rahmat, J., Dong, T., Huang, X., Tomoyasu, Y., and Halfon, M.S. (2024). Regulatory genome annotation of 33 insect species. eLife 13.

5. Bailey, T.L., Johnson, J., Grant, C.E., and Noble, W.S. (2015). The MEME Suite. Nucleic Acids Res 43, W39–49.

6. Bonasio, R. (2012). Emerging topics in epigenetics: ants, brains, and noncoding RNAs. Ann N Y Acad Sci 1260, 14–23.

7. Bonasio, R., Zhang, G., Ye, C., Mutti, N.S., Fang, X., Qin, N., Donahue, G., Yang, P., Li, Q., Li, C., et al. (2010). Genomic comparison of the ants Camponotus floridanus and Harpegnathos saltator. Science 329, 1068–1071.

8. Bonev, B., Mendelson Cohen, N., Szabo, Q., Fritsch, L., Papadopoulos, G.L., Lubling, Y., Xu, X., Lv, X., Hugnot, J.P., Tanay, A., et al. (2017). Multiscale 3D Genome Rewiring during Mouse Neural Development. Cell 171, 557–572 e524.

9. Borrelli, E., Nestler, E.J., Allis, C.D., and Sassone-Corsi, P. (2008). Decoding the epigenetic language of neuronal plasticity. Neuron 60, 961–974.

10. Burton, J.N., Adey, A., Patwardhan, R.P., Qiu, R., Kitzman, J.O., and Shendure, J. (2013). Chromosome-scale scaffolding of de novo genome assemblies based on chromatin interactions. Nat Biotechnol 31, 1119–1125.

11. Campbell, R.R., and Wood, M.A. (2019). How the epigenome integrates information and reshapes the synapse. Nature Reviews Neuroscience 20, 133–147.

12. Carmona-Aldana, F., Yong, L.W., Reinberg, D., and Desplan, C. (2024). Phenomenon of reproductive plasticity in ants. Curr Opin Insect Sci 63, 101197.

13. Corces, M.R., Trevino, A.E., Hamilton, E.G., Greenside, P.G., Sinnott-Armstrong, N.A., Vesuna, S., Satpathy, A.T., Rubin, A.J., Montine, K.S., Wu, B., et al. (2017). An improved ATAC-seq protocol reduces background and enables interrogation of frozen tissues. Nat Methods 14, 959–962.

14. Corona, M., Libbrecht, R., and Wheeler, D.E. (2016). Molecular mechanisms of phenotypic plasticity in social insects. Curr Opin Insect Sci 13, 55–60.

15. Danecek, P., Bonfield, J.K., Liddle, J., Marshall, J., Ohan, V., Pollard, M.O., Whitwham, A., Keane, T., McCarthy, S.A., Davies, R.M., et al. (2021). Twelve years of SAMtools and BCFtools. Gigascience 10.

16. Dixon, J.R., Selvaraj, S., Yue, F., Kim, A., Li, Y., Shen, Y., Hu, M., Liu, J.S., and Ren, B. (2012). Topological domains in mammalian genomes identified by analysis of chromatin interactions. Nature 485, 376–380.

17. Dudchenko, O., Batra, S.S., Omer, A.D., Nyquist, S.K., Hoeger, M., Durand, N.C., Shamim, M.S., Machol, I., Lander, E.S., Aiden, A.P., et al. (2017). De novo assembly of the Aedes aegypti genome using Hi-C yields chromosome-length scaffolds. Science 356, 92–95.

18. Durand, N.C., Robinson, J.T., Shamim, M.S., Machol, I., Mesirov, J.P., Lander, E.S., and Aiden, E.L. (2016a). Juicebox Provides a Visualization System for Hi-C Contact Maps with Unlimited Zoom. Cell Syst 3, 99–101.

19. Durand, N.C., Shamim, M.S., Machol, I., Rao, S.S., Huntley, M.H., Lander, E.S., and Aiden, E.L. (2016b). Juicer Provides a One-Click System for Analyzing Loop-Resolution Hi-C Experiments. Cell Syst 3, 95–98.

20. EpiCypher (2021). EpiCypher® CUTANATM Direct-to-PCR CUT&Tag Protocol (EpiCypher).

21. Ernst, J., and Kellis, M. (2012). ChromHMM: automating chromatin-state discovery and characterization. Nat Methods 9, 215–216.

22. Ernst, J., and Kellis, M. (2017). Chromatin-state discovery and genome annotation with ChromHMM. Nature protocols 12, 2478–2492.

23. Frost, B., Hemberg, M., Lewis, J., and Feany, M.B. (2014). Tau promotes neurodegeneration through global chromatin relaxation. Nat Neurosci 17, 357–366.

24. Fujita, Y., Pather, S.R., Ming, G.L., and Song, H. (2022). 3D spatial genome organization in the nervous system: From development and plasticity to disease. Neuron 110, 2902–2915.

25. Gallegos, D.A., Chan, U., Chen, L.F., and West, A.E. (2018). Chromatin Regulation of Neuronal Maturation and Plasticity. Trends in neurosciences 41, 311–324.

26. Gao, Q., Xiong, Z., Larsen, R.S., Zhou, L., Zhao, J., Ding, G., Zhao, R., Liu, C., Ran, H., and Zhang, G. (2020). High-quality chromosome-level genome assembly and full-length transcriptome analysis of the pharaoh ant Monomorium pharaonis. Gigascience 9.

27. Garnier, Simon, Ross, Noam, Rudis, Robert, Camargo, Pedro A., Sciaini, Marco, et al. (2024). viridis(Lite) – Colorblind-Friendly Color Maps for R.

28. Gaskill, M.M., Gibson, T.J., Larson, E.D., and Harrison, M.M. (2021). GAF is essential for zygotic genome activation and chromatin accessibility in the early Drosophila embryo. eLife 10.

29. Gasperini, M., Hill, A.J., McFaline-Figueroa, J.L., Martin, B., Kim, S., Zhang, M.D., Jackson, D., Leith, A., Schreiber, J., Noble, W.S., et al. (2019). A Genome-wide Framework for Mapping Gene Regulation via Cellular Genetic Screens. Cell 176, 377–390 e319.

30. Gene Ontology, C., Aleksander, S.A., Balhoff, J., Carbon, S., Cherry, J.M., Drabkin, H.J., Ebert, D., Feuermann, M., Gaudet, P., Harris, N.L., et al. (2023). The Gene Ontology knowledgebase in 2023. Genetics 224.

31. Ghaninia, M., Haight, K., Berger, S.L., Reinberg, D., Zwiebel, L.J., Ray, A., and Liebig, J. (2017). Chemosensory sensitivity reflects reproductive status in the ant Harpegnathos saltator. Scientific Reports 7, 3732.

32. Ghavi-Helm, Y., Klein, F.A., Pakozdi, T., Ciglar, L., Noordermeer, D., Huber, W., and Furlong, E.E. (2014). Enhancer loops appear stable during development and are associated with paused polymerase. Nature 512, 96–100.

33. Gilbert, M.B., Glastad, K.M., Fioriti, M., Sorek, M., Scarpa, T., Purnell, F.S., Xu, D., Pino, L.K., Korotkov, A., Biashad, A., et al. (2025). Neuropeptides specify and reprogram division of labor in the leafcutter ant Atta cephalotes. Cell 188, 3974–3991 e3921.

34. Gospocic, J., Glastad, K.M., Sheng, L., Shields, E.J., Berger, S.L., and Bonasio, R. (2021). Kr-h1 maintains distinct caste-specific neurotranscriptomes in response to socially regulated hormones. Cell 184, 5807–5823 e5814.

35. Gospocic, J., Shields, E.J., Glastad, K.M., Lin, Y., Penick, C.A., Yan, H., Mikheyev, A.S., Linksvayer, T.A., Garcia, B.A., Berger, S.L., et al. (2017). The Neuropeptide Corazonin Controls Social Behavior and Caste Identity in Ants. Cell 170, 748–759 e712.

36. Grandi, F.C., Modi, H., Kampman, L., and Corces, M.R. (2022). Chromatin accessibility profiling by ATAC-seq. Nature protocols 17, 1518–1552.

37. Griffith, E.C., West, A.E., and Greenberg, M.E. (2024). Neuronal enhancers fine-tune adaptive circuit plasticity. Neuron 112, 3043–3057.

38. Gurudatta, B.V., Yang, J., Van Bortle, K., Donlin-Asp, P.G., and Corces, V.G. (2013). Dynamic changes in the genomic localization of DNA replication-related element binding factor during the cell cycle. Cell Cycle 12, 1605–1615.

39. Heffel, M.G., Zhou, J., Zhang, Y., Lee, D.S., Hou, K., Pastor-Alonso, O., Abuhanna, K.D., Galasso, J., Kern, C., Tai, C.Y., et al. (2024). Temporally distinct 3D multi-omic dynamics in the developing human brain. Nature 635, 481–489.

40. Heintzman, N.D., Stuart, R.K., Hon, G., Fu, Y., Ching, C.W., Hawkins, R.D., Barrera, L.O., Van Calcar, S., Qu, C., Ching, K.A., et al. (2007). Distinct and predictive chromatin signatures of transcriptional promoters and enhancers in the human genome. Nat Genet 39, 311–318.

41. Heinz, S., Romanoski, C.E., Benner, C., and Glass, C.K. (2015). The selection and function of cell type-specific enhancers. Nat Rev Mol Cell Biol 16, 144–154.

42. Heisenberg, M. (2003). Mushroom body memoir: from maps to models. Nat Rev Neurosci 4, 266–275.

43. Hinrichs, A.S., Karolchik, D., Baertsch, R., Barber, G.P., Bejerano, G., Clawson, H., Diekhans, M., Furey, T.S., Harte, R.A., Hsu, F., et al. (2006). The UCSC Genome Browser Database: update 2006. Nucleic Acids Res 34, D590–598.

44. Hou, C., Li, L., Qin, Z.S., and Corces, V.G. (2012). Gene density, transcription, and insulators contribute to the partition of the Drosophila genome into physical domains. Mol Cell 48, 471–484.

45. Hsieh, T.H., Weiner, A., Lajoie, B., Dekker, J., Friedman, N., and Rando, O.J. (2015). Mapping Nucleosome Resolution Chromosome Folding in Yeast by Micro-C. Cell 162, 108–119.

46. Hsieh, T.S., Cattoglio, C., Slobodyanyuk, E., Hansen, A.S., Rando, O.J., Tjian, R., and Darzacq, X. (2020). Resolving the 3D Landscape of Transcription-Linked Mammalian Chromatin Folding. Mol Cell 78, 539–553 e538.

47. Hsieh, T.S., Fudenberg, G., Goloborodko, A., and Rando, O.J. (2016). Micro-C XL: assaying chromosome conformation from the nucleosome to the entire genome. Nat Methods 13, 1009–1011.

48. Hwang, J.Y., Aromolaran, K.A., and Zukin, R.S. (2017). The emerging field of epigenetics in neurodegeneration and neuroprotection. Nat Rev Neurosci 18, 347–361.

49. Ing-Simmons, E., Vaid, R., Bing, X.Y., Levine, M., Mannervik, M., and Vaquerizas, J.M. (2021). Independence of chromatin conformation and gene regulation during Drosophila dorsoventral patterning. Nat Genet 53, 487–499.

50. Institute, B. (2019). Picard toolkit (Broad Institute).

51. Jin, M.J., Wang, Z.L., Wu, Z.H., He, X.J., Zhang, Y., Huang, Q., Zhang, L.Z., Wu, X.B., Yan, W.Y., and Zeng, Z.J. (2023). Phenotypic dimorphism between honeybee queen and worker is regulated by complicated epigenetic modifications. iScience 26, 106308.

52. Jones, B.M., Waugh, A.H., Catto, M.A., Kay, S., Glastad, K.M., Goodisman, M.A.D., Kocher, S.D., and Hunt, B.G. (2025). The Fire Ant Social Chromosome Exerts a Major Influence on Genome Regulation. Mol Biol Evol 42.

53. Jones, B.M., Webb, A.E., Geib, S.M., Sim, S., Schweizer, R.M., Branstetter, M.G., Evans, J.D., and Kocher, S.D. (2024). Repeated Shifts in Sociality Are Associated With Fine-tuning of Highly Conserved and Lineage-Specific Enhancers in a Socially Flexible Bee. Mol Biol Evol 41.

54. Kaya-Okur, H.S., Janssens, D.H., Henikoff, J.G., Ahmad, K., and Henikoff, S. (2020). Efficient low-cost chromatin profiling with CUT&Tag. Nature protocols 15, 3264–3283.

55. Kaya-Okur, H.S., Wu, S.J., Codomo, C.A., Pledger, E.S., Bryson, T.D., Henikoff, J.G., Ahmad, K., and Henikoff, S. (2019). CUT&Tag for efficient epigenomic profiling of small samples and single cells. Nature communications 10, 1930.

56. Keranen, S.V.E., Villahoz-Baleta, A., Bruno, A.E., and Halfon, M.S. (2022). REDfly: An Integrated Knowledgebase for Insect Regulatory Genomics. Insects 13.

57. Kim, I.V., Navarrete, C., Grau-Bove, X., Iglesias, M., Elek, A., Zolotarov, G., Bykov, N.S., Montgomery, S.A., Ksiezopolska, E., Canas-Armenteros, D., et al. (2025). Chromatin loops are an ancestral hallmark of the animal regulatory genome. Nature 642, 1097–1105.

58. Kim, S., and Shendure, J. (2019). Mechanisms of Interplay between Transcription Factors and the 3D Genome. Mol Cell 76, 306–319.

59. Kim, S., and Wysocka, J. (2023). Deciphering the multi-scale, quantitative cis-regulatory code. Mol Cell 83, 373–392.

60. Kolde, R. (2025). pheatmap: Pretty Heatmaps.

61. Krietenstein, N., Abraham, S., Venev, S.V., Abdennur, N., Gibcus, J., Hsieh, T.S., Parsi, K.M., Yang, L., Maehr, R., Mirny, L.A., et al. (2020). Ultrastructural Details of Mammalian Chromosome Architecture. Mol Cell 78, 554–565 e557.

62. Krueger, F. (2024). TrimGalore: A wrapper tool for quality control and trimming of high-throughput sequencing data.

63. Kuang, M., Antonova, A., Khan, S., Moreno-Medina, S., and Bonasio, R. (2026a). Profiling histone modifications in Harpegnathos saltator brains by CUT&Tag [submitted].

64. Kuang, M., Shields, E.J., Khan, S., Moreno-Medina, S., and Bonasio, R. (2026b). Mapping the three-dimensional chromatin structure of the Harpegnathos saltator brain by Micro-C [submitted].

65. Langmead, B., and Salzberg, S.L. (2012). Fast gapped-read alignment with Bowtie 2. Nat Methods 9, 357–359.

66. Lawrence, M., Huber, W., Pages, H., Aboyoun, P., Carlson, M., Gentleman, R., Morgan, M.T., and Carey, V.J. (2013). Software for computing and annotating genomic ranges. PLoS computational biology 9, e1003118.

67. Lee, B.H., Wu, Z., and Rhie, S.K. (2022). Characterizing chromatin interactions of regulatory elements and nucleosome positions, using Hi-C, Micro-C, and promoter capture Micro-C. Epigenetics Chromatin 15, 41.

68. Lee, J.H., Kim, B.J., Han, G., Frunze, O., Nah, G., and Kwon, H.W. (2025). Chromosome level de Novo hybrid assembly of Asian honeybee, Apis cerana Koreana. Sci Rep 15, 26912.

69. Lettice, L.A., Heaney, S.J., Purdie, L.A., Li, L., de Beer, P., Oostra, B.A., Goode, D., Elgar, G., Hill, R.E., and de Graaff, E. (2003). A long-range Shh enhancer regulates expression in the developing limb and fin and is associated with preaxial polydactyly. Human molecular genetics 12, 1725–1735.

70. Li, H., and Durbin, R. (2010). Fast and accurate long-read alignment with Burrows-Wheeler transform. Bioinformatics 26, 589–595.

71. Li, X., Tang, X., Bing, X., Catalano, C., Li, T., Dolsten, G., Wu, C., and Levine, M. (2023). GAGA-associated factor fosters loop formation in the Drosophila genome. Mol Cell 83, 1519–1526 e1514.

72. Libbrecht, R., Oxley, P.R., Kronauer, D.J., and Keller, L. (2013). Ant genomics sheds light on the molecular regulation of social organization. Genome Biol 14, 212.

73. Love, M.I., Huber, W., and Anders, S. (2014). Moderated estimation of fold change and dispersion for RNA-seq data with DESeq2. Genome Biol 15, 550.

74. Lowe, R., Wojciechowski, M., Ellis, N., and Hurd, P.J. (2022). Chromatin accessibility-based characterisation of brain gene regulatory networks in three distinct honey bee polyphenisms. Nucleic Acids Res 50, 11550–11562.

75. Marco, A., Meharena, H.S., Dileep, V., Raju, R.M., Davila-Velderrain, J., Zhang, A.L., Adaikkan, C., Young, J.Z., Gao, F., Kellis, M., et al. (2020). Mapping the epigenomic and transcriptomic interplay during memory formation and recall in the hippocampal engram ensemble. Nat Neurosci 23, 1606–1617.

76. Martin, M. (2011). Cutadapt removes adapter sequences from high-throughput sequencing reads. EMBnetjournal 17, 3.

77. McClard, C.K., Kochukov, M.Y., Herman, I., Liu, Z., Eblimit, A., Moayedi, Y., Ortiz-Guzman, J., Colchado, D., Pekarek, B., Panneerselvam, S., et al. (2018). POU6f1 Mediates Neuropeptide-Dependent Plasticity in the Adult Brain. J Neurosci 38, 1443–1461.

78. McKenzie, S.K., and Kronauer, D.J.C. (2018). The genomic architecture and molecular evolution of ant odorant receptors. Genome Res 28, 1757–1765.

79. Merrell, A.J., and Stanger, B.Z. (2016). Adult cell plasticity in vivo: de-differentiation and transdifferentiation are back in style. Nat Rev Mol Cell Biol 17, 413–425.

80. Mohana, G., Dorier, J., Li, X., Mouginot, M., Smith, R.C., Malek, H., Leleu, M., Rodriguez, D., Khadka, J., Rosa, P., et al. (2023). Chromosome-level organization of the regulatory genome in the Drosophila nervous system. Cell 186, 3826–3844 e3826.

81. Moreno-Medina, S., Kuang, M., Liebig, J., Bonasio, R., and Khan, S. (2026a). Caste transition, behavioral monitoring, and tissue harvesting in Harpegnathos saltator [submitted].

82. Moreno-Medina, S., Kuang, M., Liebig, J., Bonasio, R., and Khan, S. (2026b). Maintaining Harpegnathos saltator colonies in the laboratory [submitted].

83. Nativio, R., Donahue, G., Berson, A., Lan, Y., Amlie-Wolf, A., Tuzer, F., Toledo, J.B., Gosai, S.J., Gregory, B.D., Torres, C., et al. (2018). Dysregulation of the epigenetic landscape of normal aging in Alzheimer’s disease. Nat Neurosci 21, 497–505.

84. Nora, E.P., Lajoie, B.R., Schulz, E.G., Giorgetti, L., Okamoto, I., Servant, N., Piolot, T., van Berkum, N.L., Meisig, J., Sedat, J., et al. (2012). Spatial partitioning of the regulatory landscape of the X-inactivation centre. Nature 485, 381–385.

85. Okamoto, N., and Yamanaka, N. (2020). Steroid Hormone Entry into the Brain Requires a Membrane Transporter in Drosophila. Curr Biol 30, 359–366 e353.

86. Opachaloemphan, C., Mancini, G., Konstantinides, N., Parikh, A., Mlejnek, J., Yan, H., Reinberg, D., and Desplan, C. (2021). Early behavioral and molecular events leading to caste switching in the ant Harpegnathos. Genes Dev 35, 410–424.

87. Opachaloemphan, C., Yan, H., Leibholz, A., Desplan, C., and Reinberg, D. (2018). Recent Advances in Behavioral (Epi)Genetics in Eusocial Insects. Annual Review of Genetics 52, 489–510.

88. Open2C, Abdennur, N., Abraham, S., Fudenberg, G., Flyamer, I.M., Galitsyna, A.A., Goloborodko, A., Imakaev, M., Oksuz, B.A., Venev, S.V., et al. (2024). Cooltools: Enabling high-resolution Hi-C analysis in Python. PLoS computational biology 20, e1012067.

89. Open2C, Abdennur, N., Fudenberg, G., Flyamer, I.M., Galitsyna, A.A., Goloborodko, A., Imakaev, M., and Venev, S.V. (2023). Pairtools: from sequencing data to chromosome contacts. bioRxiv.

90. Park, S., Alfa, R.W., Topper, S.M., Kim, G.E., Kockel, L., and Kim, S.K. (2014). A genetic strategy to measure circulating Drosophila insulin reveals genes regulating insulin production and secretion. PLoS Genet 10, e1004555.

91. Peeters, C. (1991). The occurrence of sexual reproduction among ant workers. Biological Journal of the Linnean Society 44, 141–152.

92. Peeters, C., Choe, J.C., and Crespi, B.J. (1997). Morphologically ‘primitive’ ants: comparative review of social characters, and the importance of queen–worker dimorphism. In The Evolution of Social Behavior in Insects and Arachnids, J.C. Choe, and B.J. Crespi, eds. (Cambridge: Cambridge University Press), pp. 372–391.

93. Peeters, C., and Holldobler, B. (1995). Reproductive cooperation between queens and their mated workers: the complex life history of an ant with a valuable nest. Proc Natl Acad Sci U S A 92, 10977–10979.

94. Peeters, C., Liebig, J., and Hölldobler, B. (2000). Sexual reproduction by both queens and workers in the ponerine ant Harpegnathos saltator. Insectes Sociaux 47, 325–332.

95. Penick, C.A., Ghaninia, M., Haight, K.L., Opachaloemphan, C., Yan, H., Reinberg, D., and Liebig, J. (2021). Reversible plasticity in brain size, behaviour and physiology characterizes caste transitions in a socially flexible ant (Harpegnathos saltator). Proc Biol Sci 288, 20210141.

96. Picelli, S., Bjorklund, A.K., Reinius, B., Sagasser, S., Winberg, G., and Sandberg, R. (2014). Tn5 transposase and tagmentation procedures for massively scaled sequencing projects. Genome Res 24, 2033–2040.

97. Rahman, S., Dong, P., Apontes, P., Fernando, M.B., Kosoy, R., Townsley, K.G., Girdhar, K., Bendl, J., Shao, Z., Misir, R., et al. (2023). Lineage specific 3D genome structure in the adult human brain and neurodevelopmental changes in the chromatin interactome. Nucleic Acids Res 51, 11142–11161.

98. Ramirez, F., Bhardwaj, V., Arrigoni, L., Lam, K.C., Gruning, B.A., Villaveces, J., Habermann, B., Akhtar, A., and Manke, T. (2018). High-resolution TADs reveal DNA sequences underlying genome organization in flies. Nature communications 9, 189.

99. Ramirez, F., Ryan, D.P., Gruning, B., Bhardwaj, V., Kilpert, F., Richter, A.S., Heyne, S., Dundar, F., and Manke, T. (2016). deepTools2: a next generation web server for deep-sequencing data analysis. Nucleic Acids Res 44, W160–165.

100. Rickels, R., Herz, H.M., Sze, C.C., Cao, K., Morgan, M.A., Collings, C.K., Gause, M., Takahashi, Y.H., Wang, L., Rendleman, E.J., et al. (2017). Histone H3K4 monomethylation catalyzed by Trr and mammalian COMPASS-like proteins at enhancers is dispensable for development and viability. Nat Genet 49, 1647–1653.

101. Roadmap Epigenomics, C., Kundaje, A., Meuleman, W., Ernst, J., Bilenky, M., Yen, A., Heravi-Moussavi, A., Kheradpour, P., Zhang, Z., Wang, J., et al. (2015). Integrative analysis of 111 reference human epigenomes. Nature 518, 317–330.

102. Roayaei Ardakany, A., Gezer, H.T., Lonardi, S., and Ay, F. (2020). Mustache: multi-scale detection of chromatin loops from Hi-C and Micro-C maps using scale-space representation. Genome Biol 21, 256.

103. Robinson, J.T., Thorvaldsdottir, H., Turner, D., and Mesirov, J.P. (2023). igv.js: an embeddable JavaScript implementation of the Integrative Genomics Viewer (IGV). Bioinformatics 39.

104. Robison, A.J., and Nestler, E.J. (2011). Transcriptional and epigenetic mechanisms of addiction. Nat Rev Neurosci 12, 623–637.

105. Sexton, T., Yaffe, E., Kenigsberg, E., Bantignies, F., Leblanc, B., Hoichman, M., Parrinello, H., Tanay, A., and Cavalli, G. (2012). Three-dimensional folding and functional organization principles of the Drosophila genome. Cell 148, 458–472.

106. Shen, Y., Yue, F., McCleary, D.F., Ye, Z., Edsall, L., Kuan, S., Wagner, U., Dixon, J., Lee, L., Lobanenkov, V.V., et al. (2012). A map of the cis-regulatory sequences in the mouse genome. Nature 488, 116–120.

107. Sheng, L., Shields, E.J., Gospocic, J., Glastad, K.M., Ratchasanmuang, P., Berger, S.L., Raj, A., Little, S., and Bonasio, R. (2020). Social reprogramming in ants induces longevity-associated glia remodeling. Sci Adv 6, eaba9869.

108. Shields, E.J., Sheng, L., Weiner, A.K., Garcia, B.A., and Bonasio, R. (2018). High-Quality Genome Assemblies Reveal Long Non-coding RNAs Expressed in Ant Brains. Cell reports 23, 3078–3090.

109. Shields, E.J., Sorida, M., Sheng, L., Sieriebriennikov, B., Ding, L., and Bonasio, R. (2021). Genome annotation with long RNA reads reveals new patterns of gene expression and improves single-cell analyses in an ant brain. BMC Biol 19, 254.

110. Sieriebriennikov, B., Kolumba, O., de Beaurepaire, A., Wu, J., Fambri, V., Bardol, E., Zhong, Y., Gainetdinov, I., Reinberg, D., Yan, H., et al. (2025). Transcriptional interferences ensure one olfactory receptor per ant neuron. Nature 648, 418–426.

111. Simola, D.F., Ye, C., Mutti, N.S., Dolezal, K., Bonasio, R., Liebig, J., Reinberg, D., and Berger, S.L. (2013). A chromatin link to caste identity in the carpenter ant Camponotus floridanus. Genome Res 23, 486–496.

112. Slobodyanyuk, E., Cattoglio, C., and Hsieh, T.S. (2022). Mapping Mammalian 3D Genomes by Micro-C. Methods Mol Biol 2532, 51–71.

113. Song, M., Pebworth, M.P., Yang, X., Abnousi, A., Fan, C., Wen, J., Rosen, J.D., Choudhary, M.N.K., Cui, X., Jones, I.R., et al. (2020). Cell-type-specific 3D epigenomes in the developing human cortex. Nature 587, 644–649.

114. Stadhouders, R., Thongjuea, S., Andrieu-Soler, C., Palstra, R.J., Bryne, J.C., van den Heuvel, A., Stevens, M., de Boer, E., Kockx, C., van der Sloot, A., et al. (2012). Dynamic long-range chromatin interactions control Myb proto-oncogene transcription during erythroid development. EMBO J 31, 986–999.

115. Sultan, S.E. (2021). Phenotypic Plasticity as an Intrinsic Property of Organisms. In (CRC Press).

116. Sun, C., Huang, J., Wang, Y., Zhao, X., Su, L., Thomas, G.W.C., Zhao, M., Zhang, X., Jungreis, I., Kellis, M., et al. (2021). Genus-Wide Characterization of Bumblebee Genomes Provides Insights into Their Evolution and Variation in Ecological and Behavioral Traits. Mol Biol Evol 38, 486–501.

117. Sun, L., Zhou, J., Xu, X., Liu, Y., Ma, N., Liu, Y., Nie, W., Zou, L., Deng, X.W., and He, H. (2024). Mapping nucleosome-resolution chromatin organization and enhancer-promoter loops in plants using Micro-C-XL. Nature communications 15, 35.

118. Wahl, N., Espeso-Gil, S., Chietera, P., Nagel, A., Laighneach, A., Morris, D.W., Rajarajan, P., Akbarian, S., Dechant, G., and Apostolova, G. (2024). SATB2 organizes the 3D genome architecture of cognition in cortical neurons. Mol Cell 84, 621–639 e629.

119. Wallberg, A., Bunikis, I., Pettersson, O.V., Mosbech, M.B., Childers, A.K., Evans, J.D., Mikheyev, A.S., Robertson, H.M., Robinson, G.E., and Webster, M.T. (2019). A hybrid de novo genome assembly of the honeybee, Apis mellifera, with chromosome-length scaffolds. BMC Genomics 20, 275.

120. Wang, C., Liu, C., Roqueiro, D., Grimm, D., Schwab, R., Becker, C., Lanz, C., and Weigel, D. (2015). Genome-wide analysis of local chromatin packing in Arabidopsis thaliana. Genome Res 25, 246–256.

121. Wang, M., Liu, Y., Wen, T., Liu, W., Gao, Q., Zhao, J., Xiong, Z., Wang, Z., Jiang, W., Yu, Y., et al. (2020). Chromatin accessibility and transcriptome landscapes of Monomorium pharaonis brain. Sci Data 7, 217.

122. Wickham, H. (2016). ggplot2: Elegant Graphics for Data Analysis (Springer-Verlag New York).

123. Wolff, J., Bhardwaj, V., Nothjunge, S., Richard, G., Renschler, G., Gilsbach, R., Manke, T., Backofen, R., Ramirez, F., and Gruning, B.A. (2018). Galaxy HiCExplorer: a web server for reproducible Hi-C data analysis, quality control and visualization. Nucleic Acids Res 46, W11–W16.

124. Wolff, J., Rabbani, L., Gilsbach, R., Richard, G., Manke, T., Backofen, R., and Gruning, B.A. (2020). Galaxy HiCExplorer 3: a web server for reproducible Hi-C, capture Hi-C and single-cell Hi-C data analysis, quality control and visualization. Nucleic Acids Res 48, W177–W184.

125. Xu, W., Zhong, Q., Lin, D., Zuo, Y., Dai, J., Li, G., and Cao, G. (2021). CoolBox: a flexible toolkit for visual analysis of genomics data. BMC Bioinformatics 22, 489.

126. Yan, H., Jafari, S., Pask, G., Zhou, X., Reinberg, D., and Desplan, C. (2020). Evolution, developmental expression and function of odorant receptors in insects. J Exp Biol 223.

127. Yan, H., Opachaloemphan, C., Carmona-Aldana, F., Mancini, G., Mlejnek, J., Descostes, N., Sieriebriennikov, B., Leibholz, A., Zhou, X., Ding, L., et al. (2022). Insulin signaling in the long-lived reproductive caste of ants. Science 377, 1092–1099.

128. Yan, H., Simola, D.F., Bonasio, R., Liebig, J., Berger, S.L., and Reinberg, D. (2014). Eusocial insects as emerging models for behavioural epigenetics. Nature Reviews Genetics 15, 677–688.

129. Yang, J.H., and Hansen, A.S. (2024). Enhancer selectivity in space and time: from enhancer-promoter interactions to promoter activation. Nat Rev Mol Cell Biol 25, 574–591.

130. Zhang, Y., Liu, T., Meyer, C.A., Eeckhoute, J., Johnson, D.S., Bernstein, B.E., Nusbaum, C., Myers, R.M., Brown, M., Li, W., et al. (2008). Model-based analysis of ChIP-Seq (MACS). Genome Biol 9, R137.

